# Lumi-Map, a real-time luciferase bioluminescence screen of mutants combined with MutMap, reveals *Arabidopsis* genes involved in PAMP-triggered immunity

**DOI:** 10.1101/2020.05.11.086728

**Authors:** Hiroaki Kato, Kiyoshi Onai, Akira Abe, Motoki Shimizu, Hiroki Takagi, Chika Tateda, Hiroe Utsushi, Suthitar Singkarabanit-Ogawa, Saeko Kitakura, Erika Ono, Cyril Zipfel, Yoshitaka Takano, Masahiro Ishiura, Ryohei Terauchi

## Abstract

Plants recognize pathogen-associated molecular patterns (PAMPs) to activate PAMP-triggered immunity (PTI). However, our knowledge of PTI signaling remains limited. In this report, we introduce Lumi-Map, a high-throughput platform for identifying causative single nucleotide polymorphisms (SNPs) to studying PTI signaling components. In Lumi-Map, a transgenic reporter plant line is produced that contains a firefly *luciferase* (*LUC*) gene driven by a defense gene promoter, which generates luminescence upon PAMP treatment. The line is mutagenized and the mutants with altered luminescence patterns are screened by a high-throughput real-time bioluminescence monitoring system. Selected mutants are subjected to MutMap analysis, a whole genome sequencing (WGS)-based method of rapid mutation identification, to identify the causative SNP responsible for the luminescence pattern change. We generated nine transgenic *Arabidopsis* reporter lines expressing *LUC* gene fused to multiple promoter sequences of defense-related genes. These lines generate luminescence upon activation of FLAGELLIN-SENSING 2 (FLS2) by flg22, a PAMP derived from bacterial flagellin. We selected the *WRKY29*-promoter reporter line to identify mutants in the signaling pathway downstream of *FLS2*. After screening 24,000 ethylmethanesulfonate (EMS)-induced mutants of the reporter line, we isolated 22 mutants with altered *WRKY29* expression upon flg22 treatment (abbreviated as *awf* mutants). While five flg22-insensitive *awf* mutants harbored mutations in *FLS2* itself, Lumi-Map revealed three genes not previously associated with PTI. Lumi-Map has the potential to identify novel PAMPs and their receptors as well as signaling components downstream of the receptors.

## Introduction

To defend themselves against pathogens, plants must recognize them and activate appropriate immune responses. Initial recognition of pathogens is mediated by pattern recognition receptors (PRRs) localized at the plasma membrane that sense pathogen-associated molecular patterns (PAMPs) (Boutrot and Zipfel 2017). PAMP-triggered immunity (PTI) confers broad-range resistance against pathogens (Boutrot and Zipfel 2017). Flg22, a 22 amino acid peptide derived from bacterial flagellin, is one of the most extensively studied PAMPs in plants (Boller and Felix 2009). Flg22 is recognized by the PRR FLAGELLIN-SENSING 2 (FLS2), a leucine-rich repeat receptor kinase (Gómez-Gómez and Boller 2000). FLS2 associates with the co-receptor BRASSINOSTEROID INSENSITIVE 1-ASSOCIATED RECEPTOR KINASE 1 (BAK1) and related SOMATIC EMBRYOGENESIS RECEPTOR KINASES (SERKs) in a flg22-dependent manner (Chinchilla et al. 2007; Heese et al. 2007; Schulze et al. 2010; Sun et al. 2013). Among other receptor-like cytoplasmic kinases (RLCKs), BOTRYTIS-INDUCED KINASE 1 (BIK1) associates with FLS2 (Lu et al. 2010; Zhang et al. 2010), and activates the NADPH oxidase RESPIRATORY BURST OXIDASE HOMOLOGUE PROTEIN D (RBOHD) upon PAMP binding, which leads to the apoplastic production of reactive oxygen species (ROS) in *Arabidopsis* (Kadota et al. 2014; Li et al. 2014). PAMP-mediated ROS burst is also regulated by Ca^2+^-dependent protein kinase (CDPK)-mediated phosphorylation (Boudsocq et al. 2010; Dubiella et al. 2013). MAPK cascades involving MPK3 and MPK6 are also activated and control PTI signaling at multiple levels (Rasmussen et al. 2012). Upon PAMP perception, several transcription factors involved in defense gene regulation, including those belonging to the WRKY and ethylene responsive factor (ERF) families, are phosphorylated and activated by MAPKs (Birkenbihl et al. 2017; Rasmussen et al. 2012). However, a complete molecular picture of the signaling networks regulating the rapid reprogramming of immunity-related genes remains elusive.

Forward genetic approaches have been extensively used to identify PTI signaling components. The *elfin* (*elf18-insensitive*) mutants, impaired in ROS production after elf18 treatment, included mutants of proteins involved in endoplasmic reticulum (ER) quality control (Li et al. 2009; Nekrasov et al. 2009). Boutrot et al. (2010) isolated 21 *fin* (*flagellin-insensitive*) mutants showing alterations in flg22-induced ROS burst, identifying mutants including *fin1* (corresponding to *FLS2*), *fin2* (*BAK1*), *fin3,* a mutant of *EIN2* which encodes a central regulator of ethylene-mediated signaling (Boutrot et al. 2010), and *fin4*, a mutant in the gene encoding aspartate oxidase (Macho et al. 2012). Ranf et al. (2012) screened mutant lines of *Arabidopsis* with an aequorin reporter transgene which showed changes in calcium elevation after flg22 treatment, resulting in the isolation of 35 *cce* (*changed calcium elevation*) mutants (Ranf et al. 2012). In *Arabidopsis* seedlings, sucrose-induced flavonoid accumulation is repressed upon exposure to different PAMPs (Saijo et al. 2009; Serrano et al. 2012). Using this readout, more than 50 *psl* (*priority in sweet life*) mutants were isolated, including mutants in several genes involved in ER quality control (Lu et al. 2009; Saijo et al. 2009) as well as *psl6*, an allele of *ein2* (Tintor et al. 2013). Monaghan et al. (2014) performed a screen of mutants in the *Arabidopsis bak1-5* background (*mob*: *modifier of bak1-5*) to identify mutants that restored PAMP-triggered ROS production. This screen identified the allelic *mob1* and *mob2* mutants in CPK28 (Monaghan et al. 2014) as well as *mob6*, a mutant of the gene encoding SITE-1 PROTEASE (S1P) that controls the cleavage of the endogenous peptide RAPID ALKALINIZATION FACTOR 23 (RALF23) to regulate PTI signaling (Stegmann et al. 2017). Furthermore, Li et al. (2014) reported a screening system named *Aggie* (*A*rabidopsis *g*enes *g*overning *i*mmune gene *e*xpression), in which the promoter of *FLG22-INDUCED RECEPTOR-LIKE KINASE 1* (*FRK1*) was fused with the firefly *luciferase* gene and its induction after flg22 treatment was monitored with a luminometer. This system identified two mutants: *aggie1*, a mutant of *RNA POLYMERASE II C-TERMINAL DOMAIN PHOSPHATASE-LIKE 3* (Li et al. 2014) and *aggie2*, a mutant of *POLY(ADP-RIBOSE) GLYCOHYDROLASE 1*, revealing that protein poly ADP-ribosylation plays a role in defense gene expression (Feng et al. 2015). These previous forward genetics studies of PTI signaling all involved conventional map-based cloning or a combination of map-based cloning and WGS.

Real-time bioluminescence monitoring of an organism with a reporter transgene driven by a promoter of interest is a powerful tool suitable for large-scale analyses. In this approach, gene expression is monitored in real-time with high sensitivity and accuracy in a non-destructive manner. By taking advantage of this platform, a total of 35 *Arabidopsis* circadian rhythm mutants were isolated after screening 100,000 seedlings for expression of a *luciferase* gene fused to the promoters of *GIGANTEA* and *FLOWERING LOCUS T*, which show circadian expression. This study resulted in the successful isolation of *PHYTOCLOCK1*, an essential component of the *Arabidopsis* circadian clock (Onai et al. 2004; Onai and Ishiura 2005).

Recent developments in next-generation sequencing (NGS) technologies is accelerating the WGS-based identification of mutations underlying interesting phenotypes. MutMap is one such WGS-based technique (Abe et al. 2013). In MutMap, a mutant is crossed to the original wild-type line and propagated until F_2_ progeny are obtained. DNA from 20-30 F_2_ progeny displaying the mutant phenotype are pooled and subjected to WGS. The resulting NGS short reads are aligned to the reference genome sequence of the wild-type line. A genomic region aligned to short reads with a higher frequency of SNPs points to the position of the causative mutation responsible for the phenotype. MutMap can identify the causative SNP in a mutant in a single WGS run, allowing for rapid and low-cost mutation identification.

In this report, we demonstrate Lumi-Map, which combines automated real-time luciferase bioluminescence monitoring for the high-throughput screening of mutants with MutMap to identify causative mutations involved in PTI signaling (Fig. 1). We generated nine *Arabidopsis* reporter lines that respond to the flg22 treatment. A screen of 24,000 *Arabidopsis* mutant lines with a transgenic *LUC* gene driven by the *WRKY29* gene promoter resulted in the isolation of 22 mutants with <u>a</u>ltered *<u>W</u>RKY29* expression upon flg22 treatment (*awf* mutants). The subsequent application of MutMap enabled the identification of mutated genes responsible for five mutants potentially involved in PTI.

**Fig. 1.**
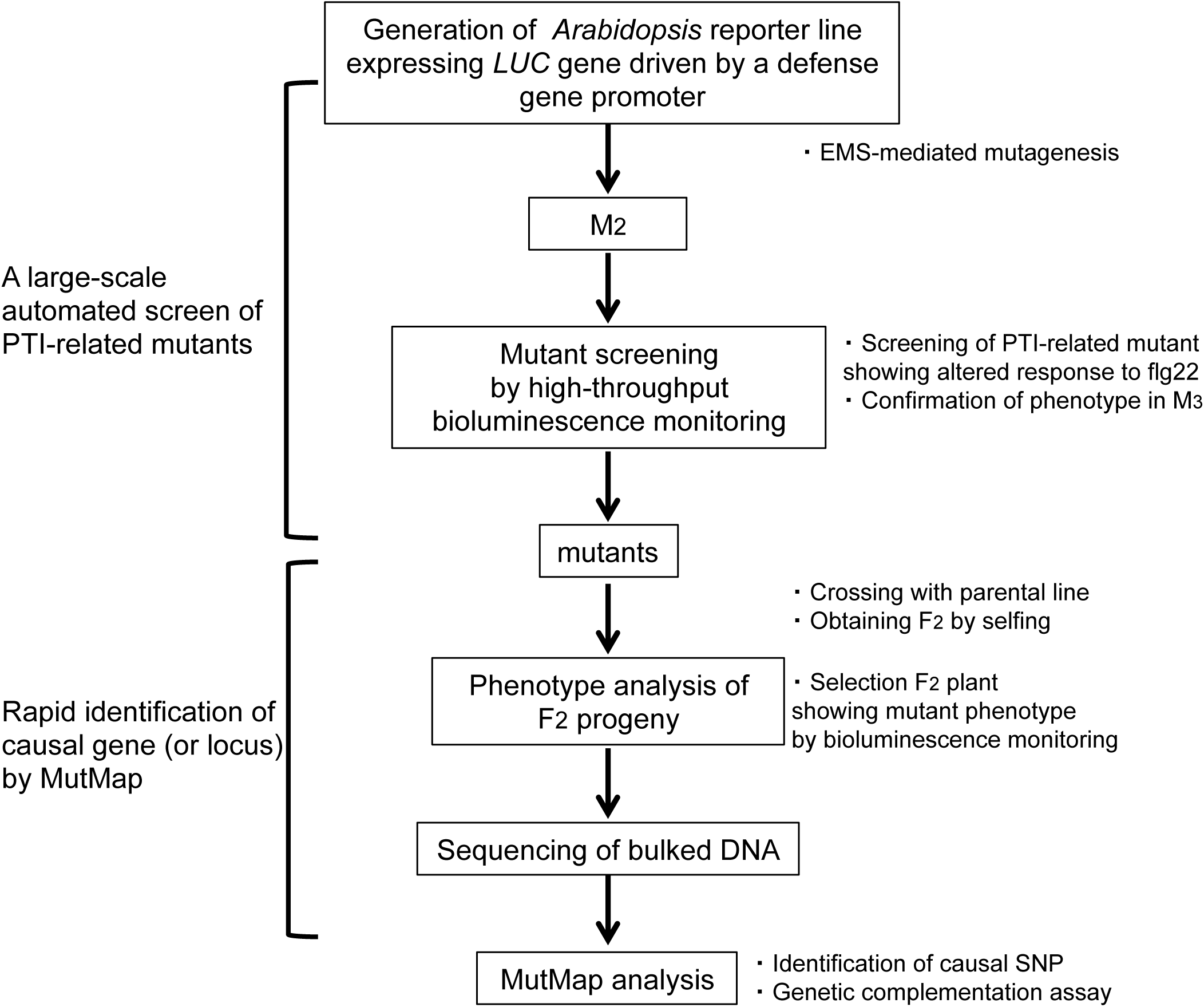
Diagram of the Lumi-Map method. A*WRKY29* reporter strain (W29-1-4) was mutagenized with ethylmethansulfonate (EMS) and M_2_ progeny were obtained. Mutant screening was performed by bioluminescence monitoring of flg22-treated M_2_ seedlings. M_2_ seedlings that showed a mutant bioluminescence phenotype were propagated to M_3_ and *awf* mutants were then chosen after confirmation of the bioluminescence phenotypes. Identification of the causal gene was performed by MutMap. *awf* mutants were crossed to the parental reporter line (W29-1-4) and F_1_ progeny were selfed to obtain the F_2_. F_2_ plants were treated with flg22 and their bioluminescence phenotypes were monitored. DNA of F_2_ plants showing mutant phenotypes were subjected to whole-genome sequencing followed by MutMap analysis to identify causal SNPs.

## Results

### Generation of transgenic *Arabidopsis* reporter lines

We generated *Arabidopsis LUC* reporter lines with promoters from nine genes (Supplementary Table S1) potentially involved in defense responses (Asai et al. 2002; Denoux et al. 2008; He et al. 2006; Navarro et al. 2006; Zipfel et al. 2004; Zipfel et al. 2006). Real-time bioluminescence monitoring revealed that these lines under liquid culture exhibited different luminescence patterns after treatment with flg22 (Fig. 2). Four lines (*WRKY29*, *WRKY18*, *WRKY28*, *RBOHD*) showed a transient induction of luminescence followed by suppression, whereas three (*PAL1*, *At1g51890*, *At2g17740*) showed an induction followed by a lasting expression, and the rest (*PR4*, *ERF1*) showed gradual increase in luminescence over the time.

**Fig. 2.**
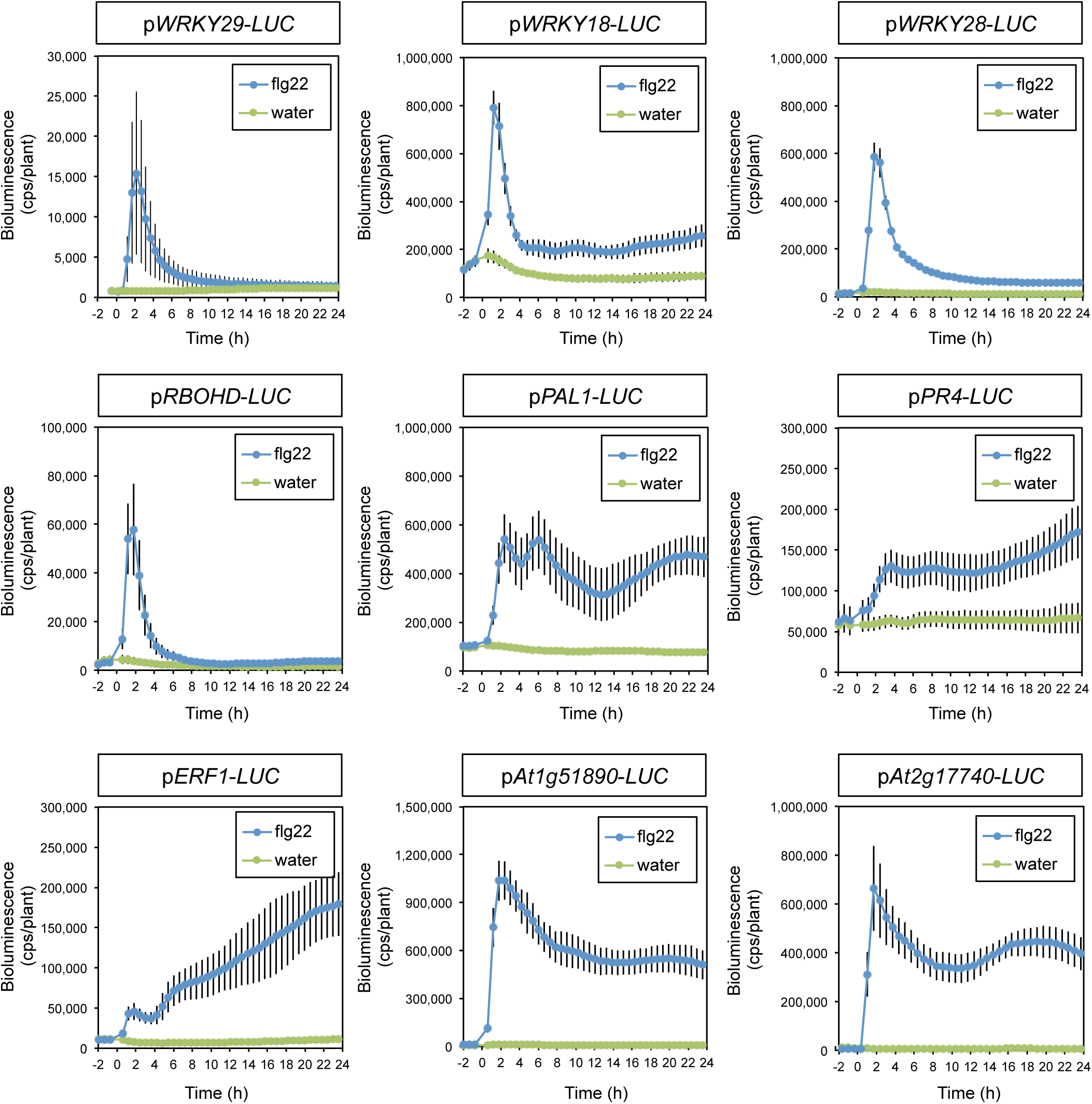
Bioluminescence patterns of nine *Arabidopsis* reporter lines after treatment with flg22. Eight day old seedlings of transgenic *Arabidopsis* plants harboring p*WRKY29*-*LUC,* p*WRKY18*-*LUC*, p*WRKY28*-*LUC*, p*RBOHD*-*LUC*, p*PAL1*-*LUC*, p*PR4*-*LUC*, p*ERF1*-*LUC*, p*At1g51890*-*LUC*, and p*At2g17740*-*LUC*, respectively, were treated with water (green) or 0.5 μM flg22 (blue). Bioluminescence from each seedling was monitored with a real-time bioluminescence monitoring system at the indicated time points. Data are shown as mean ± SE from at least seven seedlings per treatment.

### Screening *Arabidopsis* mutants affected in flg22-induced *WRKY29* expression

From among the nine reporter lines, we focused on *WRKY29* reporter line to establish Lumi-Map since previous studies indicate its role in defense response; expression of *WRKY29* is induced during flg22-triggered PTI (flg22-PTI) (Asai et al. 2002; Eulgem et al. 2000) and overexpression of *WRKY29* in *Arabidopsis* enhances resistance to *Pseudomonas syringae* pv. *maculicola* (Asai et al. 2002). *WRKY29* transcription is also known to be regulated by defense-related MAPKs such as MPK3 and MPK6 in *Arabidopsis* (Asai et al. 2002). We used *WRKY29* promoter sequence (-1931 to -1, Serrano et al. 2007) to establish a high-throughput method for monitoring gene expression during flg22-PTI. The homozygous *Arabidopsis* reporter line carrying the P*_WRKY29_*::*LUC*^+^ transgene was named W29-1-4 (Supplementary Fig. S1A), which showed a transient induction of luciferase-mediated bioluminescence under liquid culture after flg22 treatment (Supplementary Fig. S1B and S2). A similar result was obtained from the independent transgenic line W29-114A, which carried the same P*_WRKY29_*::*LUC*^+^ reporter gene, although the level of bioluminescence differed (Supplementary Fig. S1B). We also examined the level of *WRKY29* mRNA during PTI by quantitative reverse transcription-PCR (qRT-PCR) after flg22 treatment. This qRT-PCR result was overall consistent with the bioluminescence data from the W29-1-4 reporter line after treatment with flg22 (Supplementary Fig. S1C), indicating that it is possible to monitor *WRKY29* transcription levels by the bioluminescence.

To identify mutants showing altered response to flg22, we prepared EMS-mutagenized seeds of the W29-1-4 reporter line and screened 24,000 M_2_ seedlings with a high-throughput, real-time, bioluminescence monitoring system following flg22 treatment (Fig. 1). We isolated a total of 263 candidate mutants with <u>a</u>ltered *<u>W</u>RKY29* expression upon flg22 treatment (*awf* mutants) and confirmed their bioluminescence phenotypes in M_3_ progeny. We selected plants showing less than 50% or more than 200% of the maximum bioluminescence level of the original reporter line as candidate mutants. For a subset of mutant lines, M_3_ seeds could not be obtained due to abnormal growth or infertility. Mutant lines with significantly smaller seedlings than the parental line were excluded from further analysis. Finally, 22 *awf* mutants, including 18 with lower and 4 with higher bioluminescence than the wild-type, were isolated (Fig. 3, Supplementary Fig. S3, Table 1). Among the high luminescence *awf* mutants, *awf21* showed a slightly delayed induction (Fig. 3D).

**Fig. 3.**
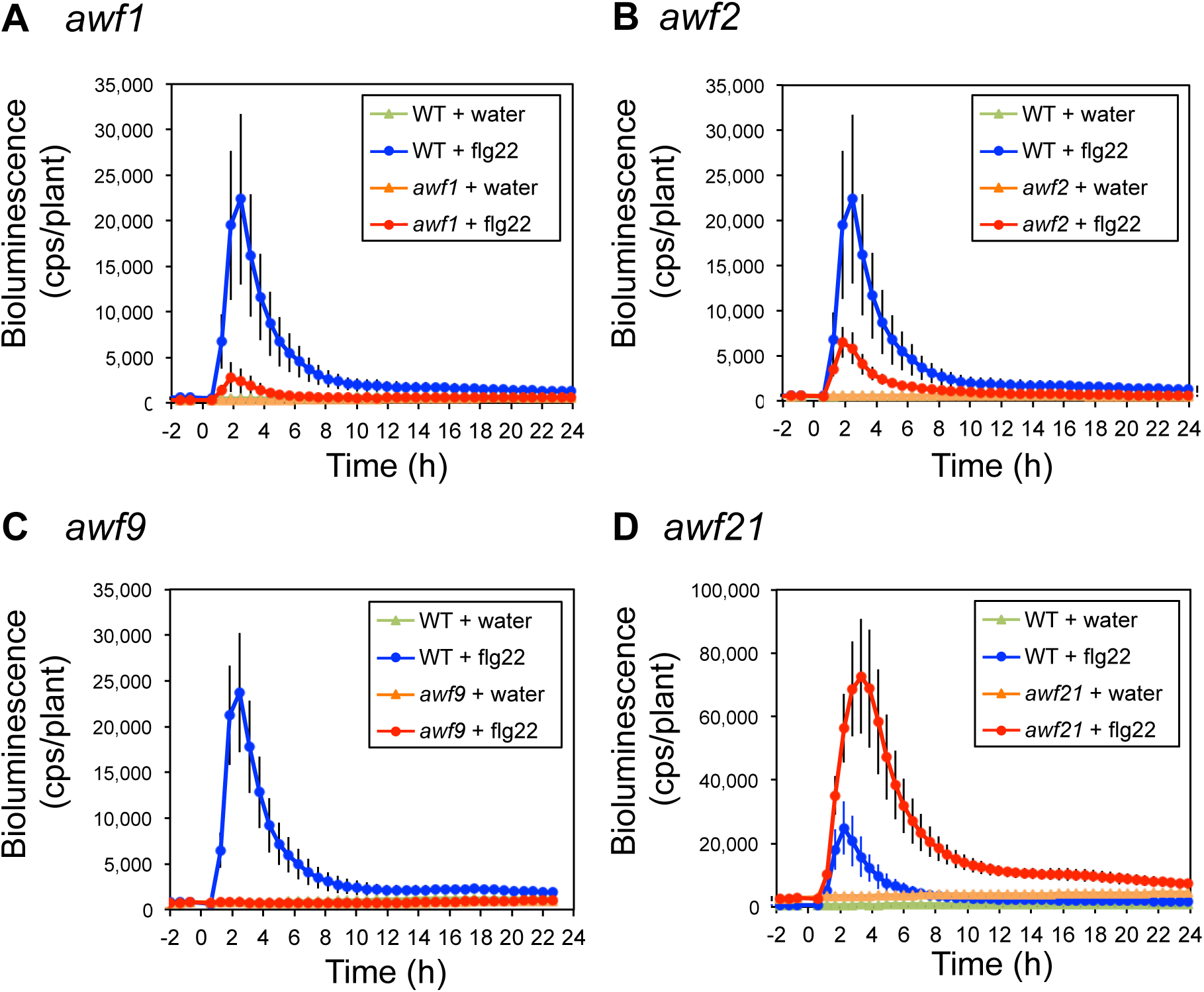
Bioluminescence patterns of mutants with altered responses to flg22 treatment. Eight day old seedlings of WT (W29-1-4) and mutants treated with water or 0.5 μM flg22. Bioluminescence from each seedling was monitored with a real-time bioluminescence monitoring system at the indicated time points. Data are shown as mean ± SE from at least 14 seedlings per treatment. Experiments were conducted three times with similar results. (A) A low bioluminescence mutant, *awf1*. (B) A low bioluminescence mutant, *awf2*. (C) A no response mutant, *awf9*. (D) A high bioluminescence and peak time altered mutant, *awf21*.

**Table 1.**
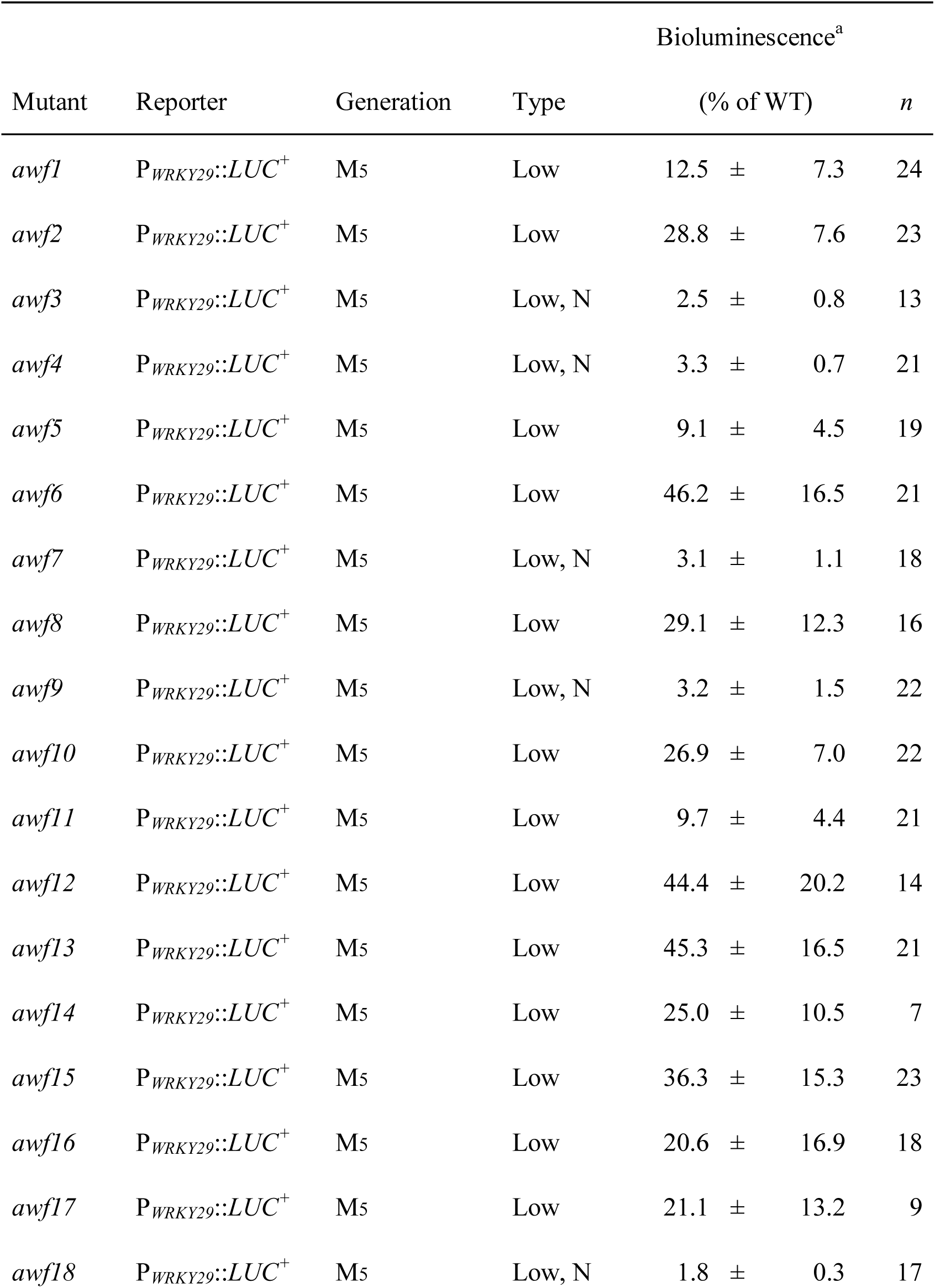
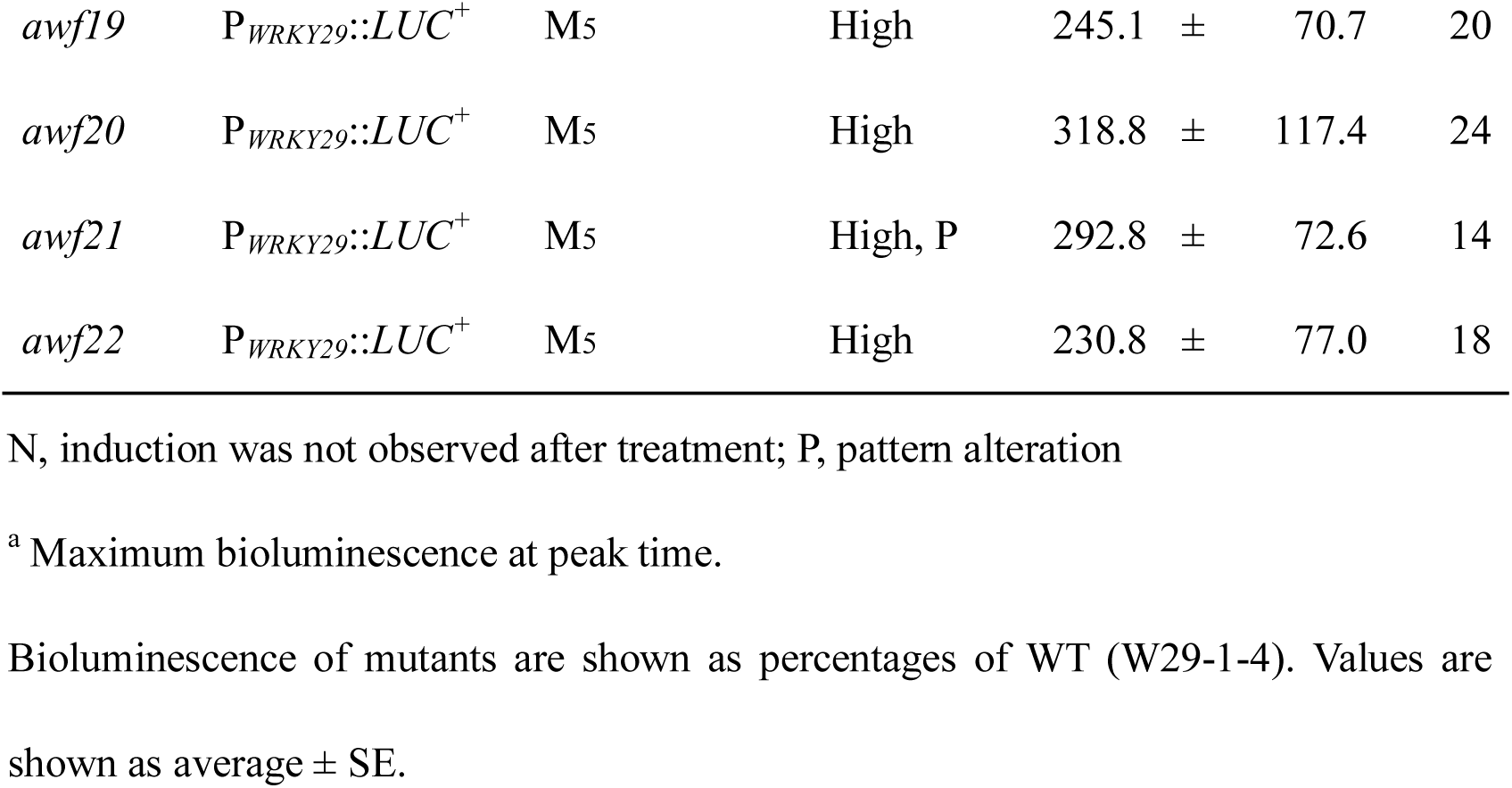
*awf* mutants and their bioluminescence phenotypes.

### Classification of the *awf* mutants

The isolated *awf* mutant lines may have mutations in the signaling pathway specific to flg22 perception by FLS2 or in a pathway shared by several PAMPs. To address this question, we studied the responses of *awf* mutants to two other PAMPs: elf18, derived from bacterial elongation factor Tu (EF-Tu), and chitin, a component of fungal cell walls. For both PAMPs, the cognate PRRs have been isolated: EF-TU RECEPTOR (EFR) for elf18 and LYSIN MOTIF RECEPTOR KINASE 5 (LYK5) acting together with CHITIN ELICITOR RECEPTOR KINASE 1 (CERK1) for chitin (Cao et al. 2014; Miya et al. 2007; Wan et al. 2008; Zipfel et al. 2006). Similar to its response to flg22 treatment, the W29-1-4 reporter line showed a transient induction of bioluminescence following elf18 and chitin treatment (Supplementary Fig. S2). We therefore treated the *awf* mutants with elf18 and chitin and grouped them based on their response to the three different PAMPs.

The low-bioluminescence mutants were classified into four groups depending on their responses to flg22, elf18, and chitin (Fig. 4A, Supplementary Fig. S4). Five mutants (*awf3*, *awf4*, *awf7*, *awf9*, and *awf18*) showed no induction of bioluminescence after flg22 treatment, while induction after elf18 and chitin treatment was not significantly altered (Group I). Three mutants (*awf11*, *awf12*, and *awf17*) showed low bioluminescence induction after flg22 treatment, while the induction by elf18 and chitin was unaltered (Group II). Three mutants (*awf6*, *awf13,* and *awf16*) showed low bioluminescence induction after both flg22 and elf18 treatment but normal induction by chitin (Group III). The remaining seven mutants (*awf1*, *awf2*, *awf5*, *awf8*, *awf10*, *awf14*, and *awf15*) showed low bioluminescence induction against all three PAMPs tested (Group IV). All four mutants with a higher bioluminescence level after flg22 treatment (*awf19*, *awf20*, *awf21*, and *awf22*) showed higher bioluminescence levels after elf18 and chitin treatments as well (Fig. 4B, Supplementary Fig. S4).

**Fig. 4.**
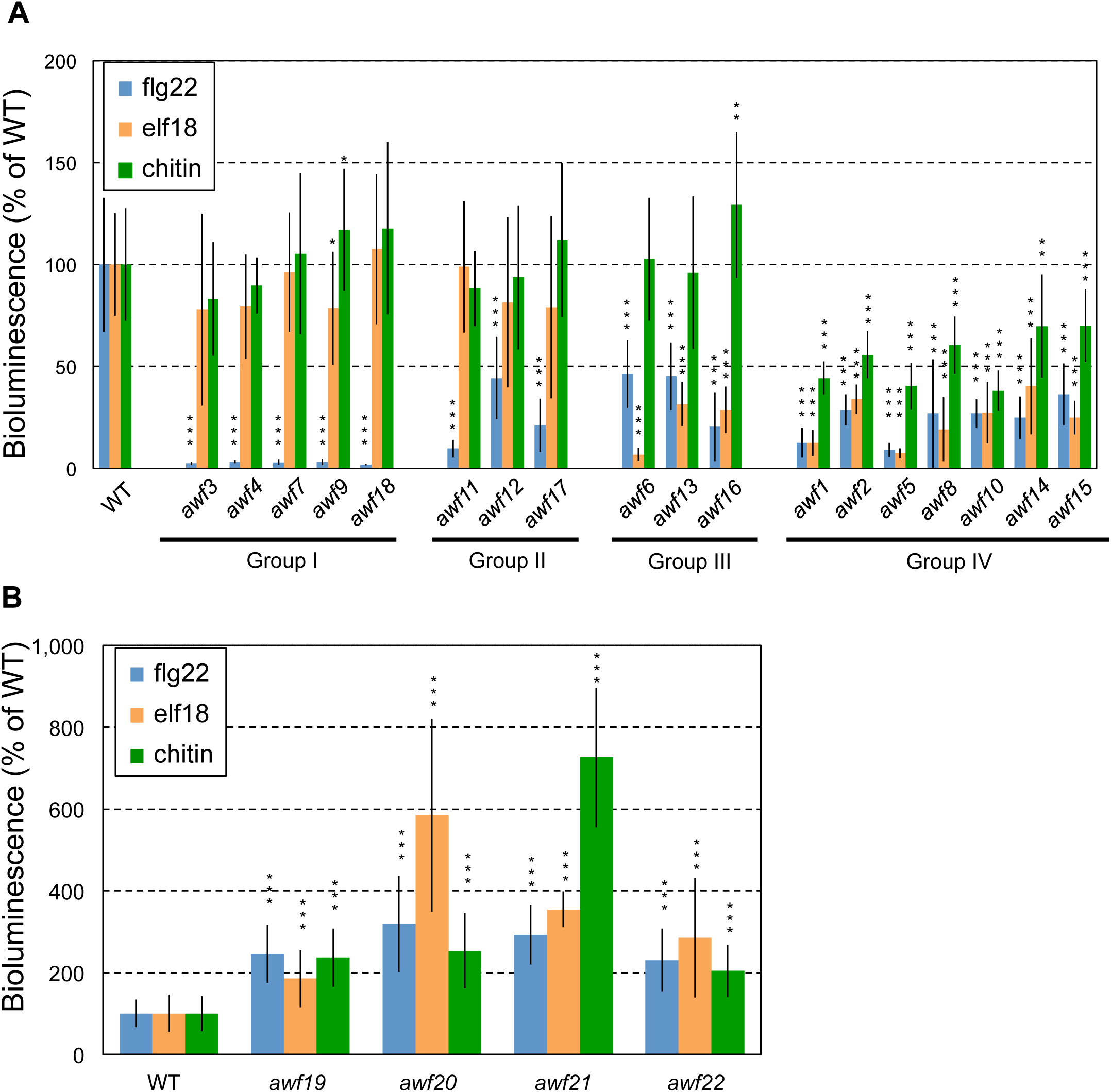
Classification of *awf* mutants by their responses to different PAMPs. Eight day old mutant seedlings were treated with 0.5 μM flg22, 0.5 μM elf18, or 1 mg/ml chitin. Response to the three elicitors was monitored with a real-time bioluminescence monitoring system. Bioluminescence is shown as % of WT (W29-1-4). Data are shown as peak mean ± SE from at least seven seedlings per treatment. (A) Low bioluminescence mutants were classified into four groups: Group I; mutants with no response to flg22, Group II; mutants showing low bioluminescence induction after flg22 treatment, Group III; mutants showing low bioluminescence induction after flg22 and elf18 treatment, Group IV; mutants showing low bioluminescence induction after flg22, elf18, and chitin treatment. (B) Mutants with increased bioluminescence induction after elicitor treatment.

### Isolation of *FLS2* mutants

FLS2 is the PRR for flg22 (Gómez-Gómez and Boller 2000). In the *fls2* mutant, PTI responses such as ROS production and MAPK phosphorylation do not occur following flg22 treatment (Asai et al. 2002; Felix et al. 1999). Five *awf* mutants showed no response after flg22 treatment (Group I, Fig. 4A, Supplementary Fig. S4), which led us to ask whether these mutants had mutations in the *FLS2* gene itself. The coding region of *FLS2* was amplified from genomic DNA from the Group I mutants (*awf3*, *awf4*, *awf7*, *awf9*, and *awf18*) by PCR and sequenced. Nucleotide substitutions were detected in the *FLS2* gene in all five Group I mutants (Fig. 5). Although they were independently isolated from different M_2_ pools, *awf3*, *awf7,* and *awf9* all shared the same mutation in the kinase domain of *FLS2* (G1042E). Further DNA sequencing of selected regions of genomes of the mutants revealed that *awf3* and *awf7* are identical presumably caused by contamination of seeds, whereas *awf9* is different from them (Supplementary Table S2). Complementation experiments confirmed that the flg22-insensitive phenotype of *awf4*, *awf9*, and *awf18* were caused by loss of FLS2 function (Supplementary Fig. S5). Thus, we conclude that mutants showing no bioluminescence activation after flg22 treatment but with unaltered responses to other PAMPs are *fls2* mutant alleles. The identification of *FLS2* mutants in our genetic screen suggests that the real-time bioluminescence monitoring platform allows uncovering PTI-related genes, including those encoding PRRs.

**Fig. 5.**
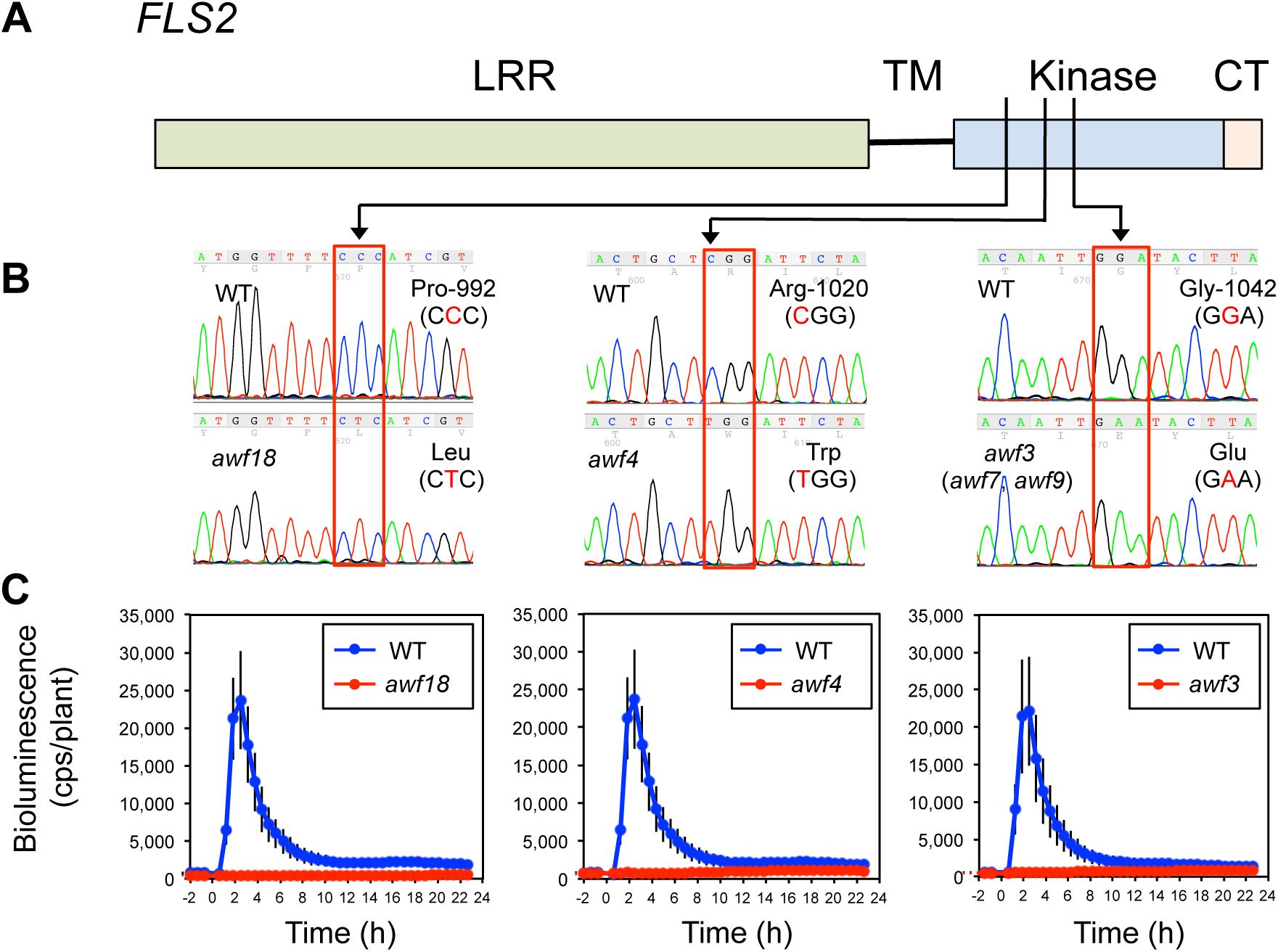
Mutations in *FLS2*. (A) Domain structure of FLS2 (LRR: leucine rich repeats; TM: transmembrane; CT:C-terminal region). (B) Confirmation of mutations in *FLS2* by Sanger sequencing. Red boxes indicate mutations in *awf18*, *awf4*, and *awf3*. *awf7* and *awf9* have the same mutation as *awf3* (G1042E). (C) Bioluminescence of mutants with mutations in *FLS2*. Eight day old seedlings of WT (W29-1-4) and mutants were treated with 0.5 μM flg22. Bioluminescence from each seedling was monitored with a real-time bioluminescence monitoring system. Data are shown as mean ± SE from at least 13 seedlings per treatment. Experiments were conducted three times with similar results.

### MutMap identifies causative genomic regions of *awf* mutants

We first applied MutMap (Abe et al. 2013) to three *awf* mutants (*awf1*, *awf2*, and *awf9*) that showed no or low bioluminescence induction after flg22 treatment. The *awf9* mutant containing a mutation in *fls2* (Fig. 5) was included as a positive control. We crossed each mutant to the W29-1-4 reporter line and obtained F_1_ seeds. The resulting F_1_ plants were self-pollinated and their F_2_ progeny were treated with flg22 and monitored for bioluminescence activation. We observed segregation of the wild-type and mutant flg22-induced bioluminescence phenotypes in an approximate 3:1 ratio (Supplementary Fig. S6, Supplementary Table S3), suggesting that all three mutants were caused by single-locus, recessive mutations. Following the MutMap procedure, we pooled DNA from 30 F_2_ individuals displaying the mutant phenotype. Equal amounts of leaf material were obtained from each individual and mixed together, and DNA was extracted from this mixture. This pooled DNA was subjected to WGS using an Illumina NGS platform. We obtained 25-30 million sequence reads from each of the three mutants (Supplementary Table S4). These reads were aligned to the W29-1-4 reference sequence and SNPs were identified. For each genomic position containing a SNP, the SNP-index (the frequency of short reads containing SNPs different from the reference) was calculated and graphs relating SNP positions to SNP-indices were generated for the five *Arabidopsis* chromosomes (Fig. 6, Supplementary Table S5). For all three mutants, we identified single genomic regions harboring a cluster of SNPs with a SNP-index > 0.95; *awf1* showed a SNP-index peak on chromosome 1, whereas *awf2* and *awf9* had SNP-index peaks on chromosome 5 (Fig. 6). The position of the SNP-index peak of *awf9* exactly corresponded to the location of *FLS2* (SNP-18794926 in chr. 5). These results demonstrate that Lumi-Map, bioluminescence monitoring combined with MutMap, rapidly and effectively identifies the position of causative mutations for mutants with altered responses to flg22 treatment.

**Fig. 6.**
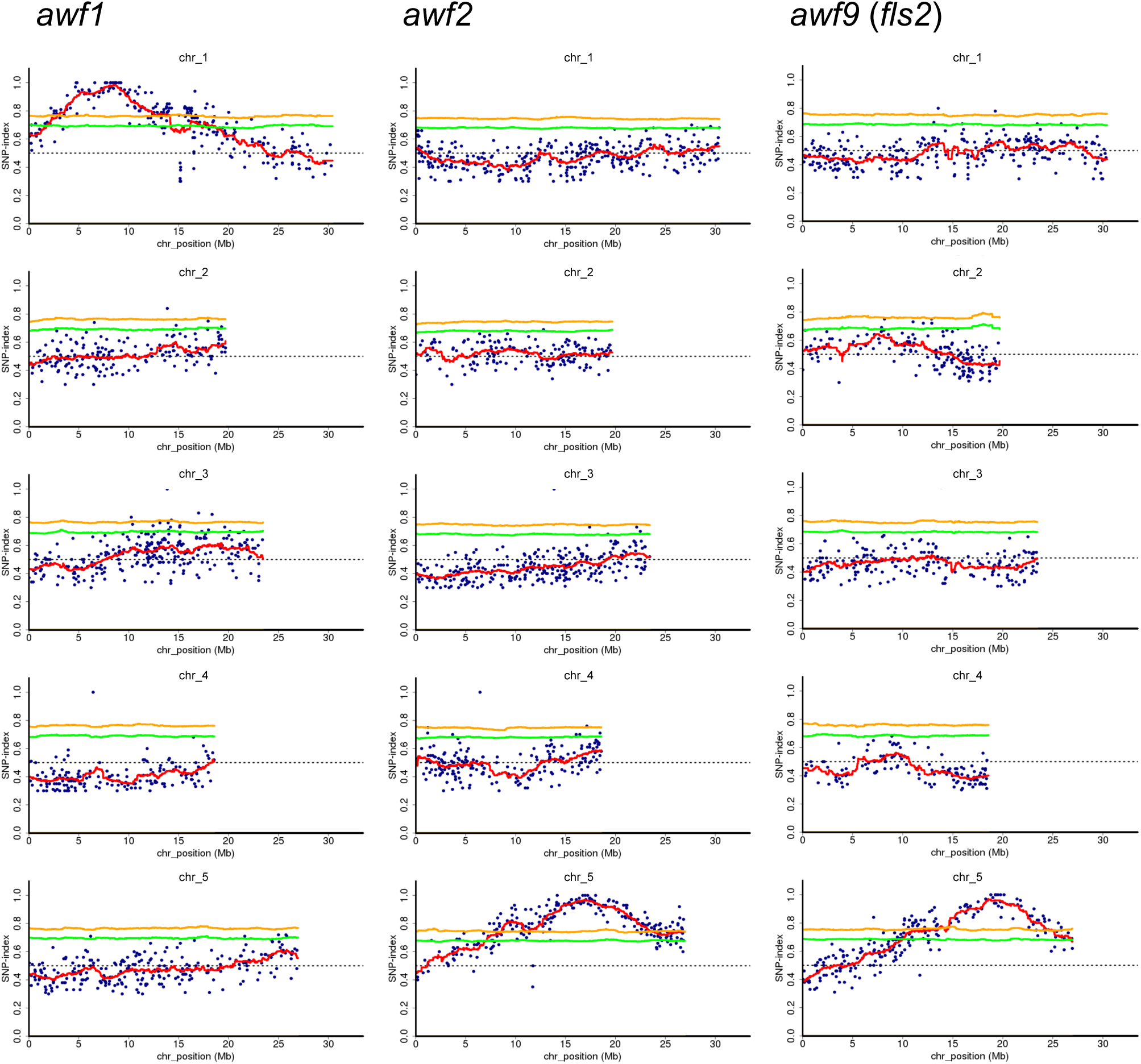
Application of MutMap to *awf1*, *awf2*, and *awf9*. SNP-index plots of the five chromosomes of *Arabidopsis* generated by MutMap analysis, showing a genomic region with the highest SNP-index peak harboring the candidate mutation. Each mutant was crossed to the parental reporter line (W29-1-4) and resulting F_2_ progeny were tested for their bioluminescence phenotype after flg22 treatment. Bulked DNA from 30 F_2_ progeny showing mutant phenotypes were used for sequencing and MutMap analysis. Blue dots correspond to SNPs identified in the mutant lines relative to W29-1-4. The red line represents average SNP-index values across a 2 Mb sliding window with 10 kb increments. The green and yellow lines show the 95% and 99% confidence limit, respectively, of SNP-index values under the null hypothesis of SNP-index = 0.5.

Among the other 11 low bioluminescence mutants that seem to be non-*fls2* mutants, we extended MutMap to four other mutants (*awf5*, *awf8*, *awf14,* and *awf16*). We observed segregation in the F_2_ (Supplementary Fig. S6, Supplementary Table S3). We then pooled DNA from 30 F_2_ mutant progeny and subjected this DNA to MutMap. In all four cases, we identified single genomic regions with SNP-index peaks: *awf5* and *awf14* showed SNP-index peaks on chromosome 5, whereas *awf8* and *awf16* had SNP-index peaks on chromosome 2 (Supplementary Fig. S7). Of the seven low bioluminescence mutants applied to MutMap, none of the loci identified by MutMap overlapped (Fig. 6, Supplementary Fig. S7), suggesting that different mutations are responsible for the observed phenotypes.

We also performed MutMap on *awf19* and *awf21*, which showed higher bioluminescence upon flg22 treatment and observed segregation in the F_2_ between plants with wild-type and higher bioluminescence levels (Supplementary Fig. S7, Supplementary Table S3). MutMap identified a single genomic region harboring a SNP-index peak on chromosome 5 for *awf19*, and chromosome 1 for *awf21* (Supplementary Fig. S7).

### Identification of the causative mutations of selected *awf* mutants

For the *awf1* mutant, we examined SNPs with a SNP-index > 0.95 in detail (Supplementary Table S6). One SNP, SNP-8074904, was a nonsense mutation located in *Ethylene responsive factor 019* (*ERF019*) (*AT1G22810*) changing a Trp residue to a stop codon (Fig. 7A, B, Supplementary Table S6). The *ERF* family in *Arabidopsis* consists of 12 groups and *ERF019* is a member of group IIc (Nakano et al. 2006). A class IIc *ERF* in rice, *SERF1*, is reported to be involved in ROS signaling during salt stress response (Schmidt et al. 2013). We therefore hypothesized that the nonsense mutation in *ERF019* underlies the *awf1* mutant. To test this hypothesis, we carried out a complementation assay by transforming the *awf1* mutant with wild-type *ERF019* under the control of its native promoter. Bioluminescence following flg22 treatment was restored in the complemented transformants (Fig. 7C, Supplementary Fig. S8A). Thus, MutMap successfully identified SNP-8074904 as the causative mutant of the *awf1* phenotype.

**Fig. 7.**
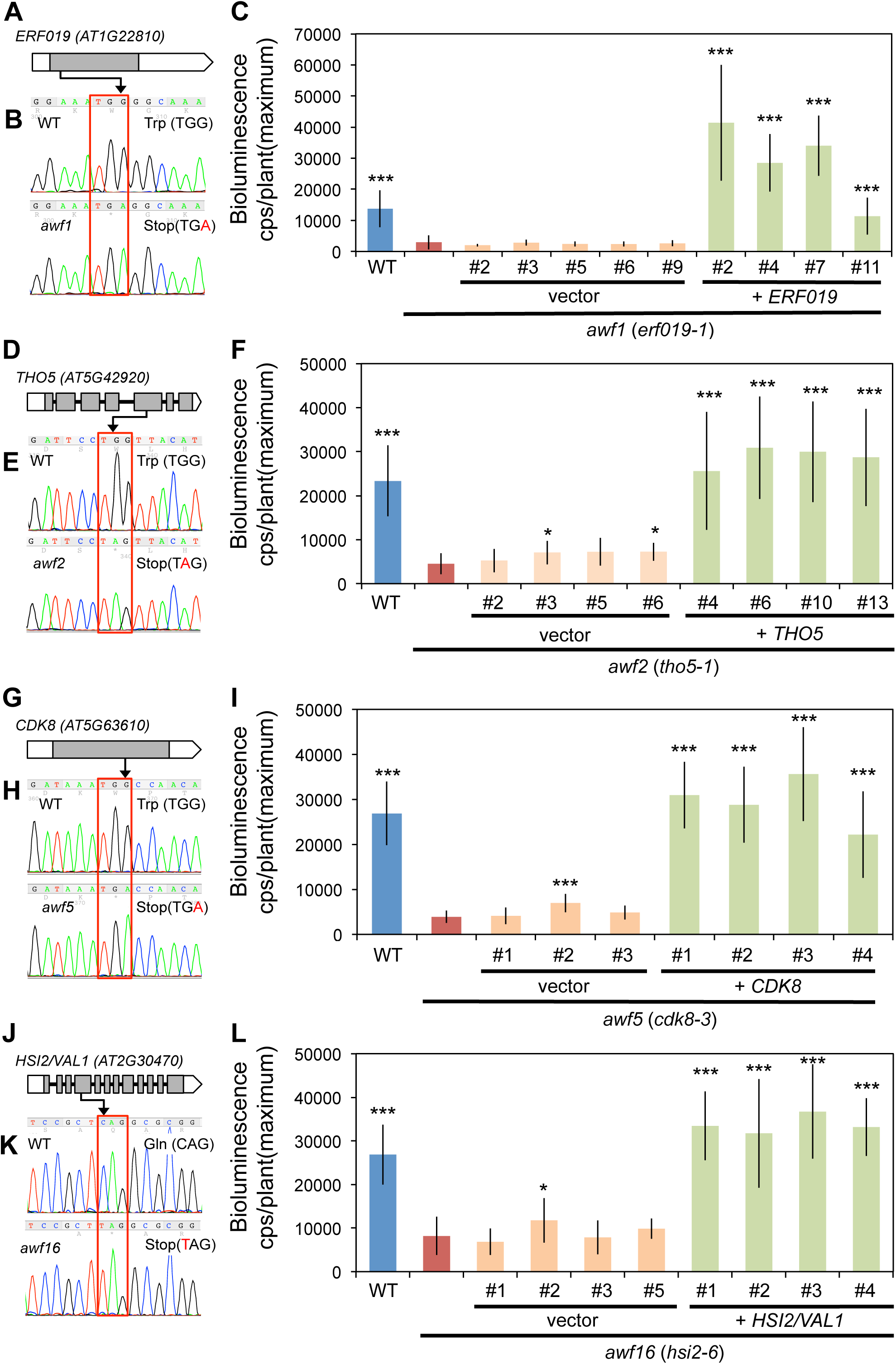
Genetic complementation of low bioluminescence mutants. Genetic complementation of *awf1* with *ERF019* (A-C), *awf2* with *THO5* (D-F), *awf5* with *CDK8* (G-I), and *awf16* with *HSI2/VAL1* (J-L). Structure of *ERF019* (*AT1G22810*) (A), *THO5* (*AT5G42920*) (D), *CDK8* (AT5G63610) (G), and *HSI2/VAL1* (*AT2G30470*) (J) are shown. Open and gray boxes represent untranslated regions (UTRs) and exons, respectively, while lines denote introns. Confirmation of the mutations by Sanger sequencing (B, E, H, and K). Red boxes indicate the mutation sites in *awf1* (B), *awf2* (E), *awf5* (H), and *awf16* (K). Eight day old seedlings of WT (W29-1-4), the four mutants *awf1*, *awf2*, *awf5*, *awf16*, as well as mutants transformed with the indicated gene or empty vector (vector) were treated with 0.5 μM flg22 and bioluminescence was monitored with a real-time bioluminescence monitoring system. Data are shown as peak mean ± SE from at least five seedlings per line (C, F, I, and L). Asterisks indicate significant differences compared with the bioluminescence of respective *awf* mutant (*; *P* < 0.05, ***; *P* < 0.001, two-tailed t-tests).

Using such complementation experiments, we also validated the causative mutation in *awf2* mutant within *THO5* (*AT5G42920*), a member of the THO/TREX (suppressors of the **t**ranscription defects of ***h****pr1Δ* mutants by **o**verexpression-**Tr**anscription and **Ex**port) protein family, (Fig. 7D to F, Supplementary Fig. S8B), *awf5* mutant within *CYCLIN-DEPENDENT KINASE 8* (*CDK8*) (*AT5G63610*) (Fig. 7G to I, Supplementary Fig. S8C), and in *awf16* mutant within *HIGH-LEVEL EXPRESSION OF SUGAR-INDUCIBLE 2* (*HSI2*) /*VIVIPAROUS1/ABA INSENSITIVE3-LIKE1* (*VAL1*) (*AT2G30470*) (Fig. 7J to L, Supplementary Fig. S8D). These results demonstrate that applying MutMap to mutants isolated by bioluminescence monitoring successfully identifies causative mutations. MutMap analysis of other *awf* mutants did not narrow down single candidate but exhibited multiple candidates so that we did not attempt complementation in this study.

### Gene expression in the selected mutants, *awf1, awf2,* and *awf16*

We selected the three mutants *awf1* (*erf019-1*), *awf2* (*tho5-1*), and *awf16* (*hsi2-6*) for further characterization. We included *awf4* (*fls2*) as a control and excluded *awf5* (*cdk8-3*) as *CDK8* has already been shown to be involved in anti-bacterial and anti-fungal defense in *Arabidopsis* (Huang et al. 2019; Zhu et al. 2014). Before proceeding, we confirmed expression of *WRKY29* gene after flg22 treatment in the mutants by qRT-PCR (Fig. 8A, Supplementary Fig. S9) at the eight-day-old seedling stage as used for bioluminescence measurement. In the *awf1* (*erf019-1*) and *awf16* (*hsi2-6*), the expression of *WRKY29* gene showed a clear reduction which is in line with the reduction in bioluminescence. However, in the *awf2* (*tho5-1*) mutant, we found the level of *WRKY29* transcript was not altered, which was inconsistent with the results of bioluminescence and qRT–PCR of *Luciferase* transcript (Figs. 3, 8A, Supplementary Fig. S9). We hypothesize that the regulatory elements interacting with the promoter sequence of *WRKY29* (-1931 to -1, Serrano et al. 2007) used for driving *LUC* was indeed affected by the *tho5* mutation. However, it is possible that additional mechanisms are involved in controlling the endogenous *WRKY29* transcript level (see Discussion).

**Fig. 8.**
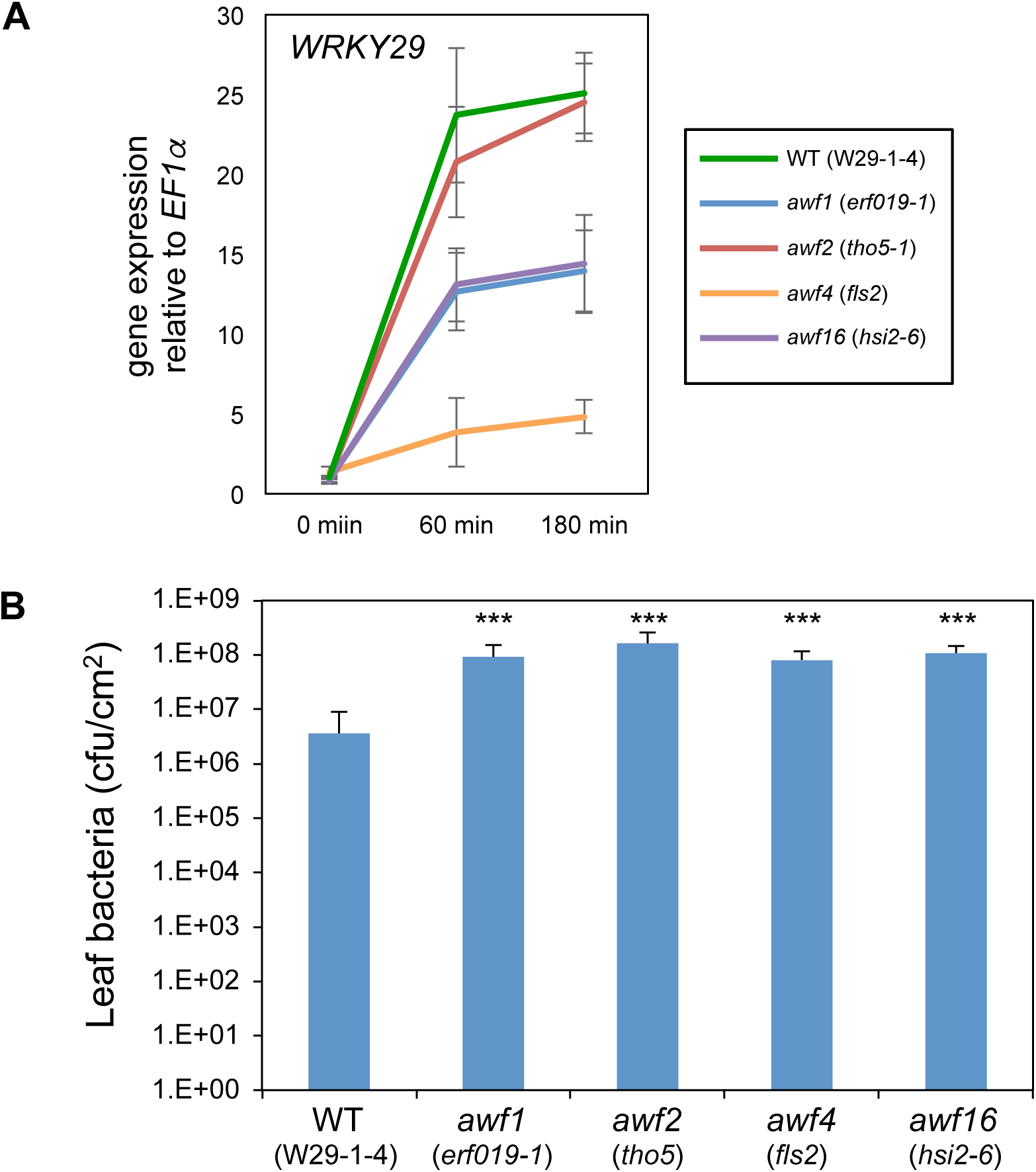
PTI analysis in *awf1, awf2,* and *awf16* mutants. (A) qRT-PCR analysis of *WRKY29* gene expression in *awf* mutants. Eight day old seedlings of WT (W29-1-4), *awf1* (*erf019-1*), *awf2* (*tho5-1*), *awf4* (*fls2*), and *awf16* (*hsi2-6*) were treated with 0.5 μM flg22. Gene expression values of *WRKY29* are relative to the *EF1α* housekeeping gene (*At1g07920*) and were normalized to untreated WT seedlings. Values are shown as mean ± SE. Experiments were conducted twice with similar results. (B) Enhanced susceptibility of *awf1, awf2,* and *awf16* mutants to *Pto DC3000*. Five-week-old plants of WT (W29-1-4), *awf1* (*erf019-1*), *awf2* (*tho5-1*), *awf4* (*fls2*), *awf16* (*hsi2-6*), as well as mutants transformed with the indicated gene were inoculated with *Pto DC3000* (inoculation dose, OD_600_ = 0.01). Bacterial titers were determined 3 days post inoculation. Values are mean ± SE (*n* = 6). Asterisks indicate significant differences compared with leaf bacteria of WT (***; *P* < 0.001, two-tailed t-tests).

### PAMP-induced apoplastic ROS production and *WRKY29* gene expression can be uncoupled

After PAMP perception by PRRs, PTI signaling branches; one pathway leads to ROS production and the other to MAPK activation (Segonzac et al. 2011; Xu et al. 2014). The *awf* mutants were isolated by screening for mutants with alterations in *WRKY29* promoter induction, which is reported to be regulated by MAPK signaling (Asai et al. 2002; Eulgem et al. 2000). To test whether these mutants also have alterations in the ROS pathway, we performed a ROS assay by treating leaves with flg22. *Awf1* (*erf019-1*), *awf2* (*tho5-1*), and *awf16* (*hsi2-6*) showed ROS generation patterns after flg22 treatment similar to wild-type Col-0 plants and the W29-1-4 reporter line (Supplementary Fig. S10). These results indicate that the mutants tested here have no alternations in the pathway leading to early ROS generation after flg22 treatment. As expected, *awf4* (*fls2*) did not show flg22-induced ROS generation.

### MAP kinase activity was not affected in *awf1, awf2,* and *awf16*

We also tested activation of MAPK after elicitation of seedlings of *awf1* (*erf019-1*), *awf2* (*tho5-1*), *awf16* (*hsi2-6*) as well as *awf4* (*fls2*) mutants after flg22 treatment (Supplementary Fig. S11). *awf1* and *awf16* mutants showed MAPK activation presumably of MPK6 (approx. 47 kDa), MPK3 (approx. 43 kDa) and MPK4 (approx. 38 kDa) (Frei dit Frey et al. 2014; Ranf et al. 2011) similar to the wild-type W29-1-4 reporter, whereas *awf4* (*fls2*) mutant showed no MAPK activation after flg22 treatment as expected (Supplementary Fig. S11).

### *ERF019, THO5,* and *HSI2* regulate plant immunity

To test whether the genes identified in this study indeed regulate *Arabidopsis* immunity, we challenged five-week-old plants of the mutants with *P. syringae* pv. *tomato* (*Pto*) DC3000. We found that *awf1* (*erf019-1*), *awf2* (*tho5-1*), and *awf16* (*hsi2*) were more susceptible to *Pto* DC3000 than wild-type W29-1-4 reporter line (Fig. 8B), and their susceptibility was comparable to that of the *awf4* (*fls2*) mutant. These results indicate that the identified genes are involved in the basal resistance against *Pto* DC3000. We also tested *WRKY29* expression after flg22 treatment in five-week-old plants by qRT-PCR (Supplementary Fig. S12). At 60 min after elicitation, *WRKY29* expression was lower in all the three *awf* mutants, whereas at 180 min after elicitation, there was no significant *WRKY29* downregulation observed for *awf2* (*tho5-1*) and *awf16* (*hsi2-6*) (Supplementary Fig. S12).

## Discussion

### Real-time bioluminescence monitoring for the study of PTI signaling

Here, we used Lumi-Map to successfully identify known and novel PTI signaling components. The major advantage of using real-time bioluminescence monitoring for gene expression studies is that it enables high-throughput analysis of gene expression kinetics in a highly sensitive and quantitative manner over longer (∼24 hours) time periods (Kondo and Ishiura 1994; Onai et al. 2004). By comparing bioluminescence induction patterns, mutants that alter the transcriptional regulation of the reporter gene can be rapidly identified. In previous reports, two *aggie* mutants were isolated using a p*FRK1*:*LUC* reporter *Arabidopsis* strain (Feng et al. 2015; Li et al. 2014). In this screen, bioluminescence was observed at only one time point after elicitation using a conventional luminometer and map-based cloning was employed to identify the causative mutations.

To identify clock genes in diverse organisms, Ishiura and colleagues have developed a series of automated devices for high-throughput real time monitoring of bioluminescence (Kondo and Ishiura 1994; Kondo et al. 1994; Okamoto et al. 2005a; Okamoto et al. 2005b; Okamoto et al. 2005c; Onai et al. 2004). Using one of these devices, they succeeded in identifying the gene *PHYTOCLOCK1* which is essential for the *Arabidopsis* circadian clock (Onai et al. 2004; Onai and Ishiura 2005), thereby demonstrating the power of this phenotyping platform. In the current study, we employed the same device (see Materials and Methods) to address PTI signaling.

### Identification of novel genes involved in PTI responses

We identified *ERF019* as a gene that regulate *WRKY29* gene expression during flg22, elf18, and chitin PTI responses (Fig. 4, Supplementary Fig. S4). This gene is also involved in basal resistance to *Pto* infection (Fig. 8). It was recently reported that *ERF019*–overexpressing *Arabidopsis* also showed increased susceptibility against *Pto* infection (Huang et al. 2019). In *Arabidopsis,* ERFs have been implicated in plant defense responses. Knockdown of some ERFs by RNAi enhanced susceptibility to *Pto* (Zhang et al. 2015; Zhang et al. 2016). ERF6 is phosphorylated by MPK3 and MPK6 during *Botrytis cinerea* infection (Meng et al. 2013). ERF104 is reported to be involved in flg22-mediated signaling pathway (Bethke et al. 2009). It is possible that *ERF019* regulates the transcription of WRKYs, including *WRKY29*, in the MAPK pathways responding to bacterial infection. The *erf019* mutant showed normal ROS production and MAPK activation after flg22 elicitation (Supplementary Figs. S10, S11), suggesting that ERF019 functions downstream of ROS signaling and the MAPK cascade.

From the mutants showing low bioluminescence after flg22 treatment, we identified *awf2* mutant with a defect in *THO5* gene (Figs 3, 6, 7). *THO5* is also involved in basal resistance against *Pto* infection (Fig. 8B). However, our qRT-PCR of endogenous *WRKY29* mRNA did not reveal downregulation after flg22 elicitation (Fig. 8A, Supplementary Fig. S9). We hypothesize that this discrepancy of the results may be explained by the promoter sequence used to drive *LUC* gene, which was a 1.9 kb fragment (-1931 to -1, Serrano et al. 2007) upstream of the *WRKY29* gene start codon. The possible regulatory elements binding to this 1.9 kb fragment may have been affected by THO5 (Fig. 3, Supplementary Fig. S9). However, the endogenous *WRKY29* gene expression may be regulated by additional *cis* elements located beyond the 1.9 kb region and interactors to such elements may have masked the effect of *tho5* that was observed with a shorter promoter used to drive *LUC.* THO5 is a member of THO/TREX family which is involved in mRNA transport and microRNA synthesis (Francisco-Mangilet et al. 2015; Sørensen et al. 2017; Yelina et al. 2010). Therefore, it could be also possible that *tho5* mutation may have altered the post-transcriptional regulation of *WRKY* transcripts. Future works are required to understand this discrepancy. This observation poses a caveat for reporter analysis that we have to use a sufficiently long promoter sequence to drive *LUC* gene to faithfully monitor the expression of the target gene.

We also identified a mutation in *HSI2*/*VAL1* as causative for the *awf16* mutant (Fig. 7). A mutation in *HSI2*/*VAL1* was previously identified as causative for an *Arabidopsis* sugar response mutant where it seems to function as a transcriptional repressor (Tsukagoshi et al. 2005). It also functions as a transcriptional repressor in flowering and drought stress response (Qüesta et al. 2016; Sharma et al. 2013), demonstrating that it plays a role in diverse processes in *Arabidopsis*. It will be interesting to study whether *HSI2*/*VAL1* impinges on flg22 and elf18 signaling pathways through transcriptional repression.

The flg22/FLS2 pathway is the best characterized PTI pathway; many genes involved in this pathway have been described. *Arabidopsis* mutants defective in ROS production following flg22 treatment have been previously studied (Boutrot et al. 2010), resulting in the isolation of the ethylene signaling protein EIN2 and aspartate oxidase (Boutrot et al. 2010; Macho et al. 2012). However, the genes identified in the current study (*ERF019, THO5,* and *HSI2/VAL1*) did not overlap with the genes previously isolated in the flg22/FLS2 pathway, and are not involved in early ROS generation nor activation of MAPK cascade after flg22 perception by FLS2 (Supplementary Figs. S10, S11). These results indicate that the genes isolated in the current study function downstream of the ROS and MAPK pathways. The reason that we did not recover *bak1* mutants in our screen may be because we have removed mutants with reduced sizes from our consideration. Most loss-of-function *bak1* mutation are indeed known to cause plant size reduction (Chinchilla et al. 2007; Wierzba and Tax 2016).

Here, we developed Lumi-Map to rapidly identifying genes potentially involved in the downstream signaling of PTI (Fig. 1). Large-scale application of Lumi-Map could enable the identification of the full repertoire of signaling components linking a cell surface receptor to a reporter gene. For this purpose, we have already generated *Arabidopsis Luciferase* reporter lines with promoters from nine genes exhibiting different response patterns to flg22 (Fig. 2) and are going to enlarge the repertoire, which will be freely available to the research community. A large number of mutated genes identified by Lumi-Map may be shared between different PAMP/PRR pairs, allowing the position of the gene in the signaling networks to be inferred. As shown by the identification of *fls2* in our screen, mutants that are non-responsive to a specific PAMP are likely mutated in the gene encoding the cognate PRR. This result suggests Lumi-Map as a potentially powerful platform to identify novel PRR-encoding genes and their downstream signaling genes using various PAMPs or microbial extracts as input signals.

## Materials and Methods

### Plant materials and growth condition

*Arabidopsis thaliana* (ecotype Col-0) seeds were used for the construction of all transgenic lines. Surface-sterilized seeds of *Arabidopsis* were incubated at 4°C in the dark for 2 days and grown on Murashige and Skoog (MS) medium (Murashige and Skoog 1962) containing 1.5% (w/v) sucrose and 0.8% (w/v) agar at 23°C in light. For seed propagation, we grew *Arabidopsis* plants on MS agar media for 10-14 days before transferring to soil at 23°C in continuous light from white fluorescent lamps at a light intensity of 57 μmol m^-2^sec^-1^.

### Molecular cloning

We confirmed the nucleotide sequences of all constructs using standard DNA manipulation and sequencing techniques. Primers used in this study are listed in Supplementary Table S7.

The P*_WRKY29_*::*LUC*^+^ reporter gene cassette consisted of the *Arabidopsis WRKY29* promoter (P*_WRKY29_*; -1931 to -1, Serrano et al. 2007), the coding region of a modified firefly *luciferase* gene (*LUC*^+^, Promega Corp, Madison, WI, USA), and T*_NOS_*. The P*_WRKY29_*::*LUC*^+^ reporter gene cassette was inserted into pBIB-HYG (Becker 1990) with *Sal*I and *Kpn*I. Reporter gene cassettes for eight other reporter lines were constructed using the promoter sequences of respective genes (Table S1).

For complementation of *awf1,* a 4.0 kb fragment containing the *ERF019* gene and its native promoter was amplified and inserted into pBI101 with *Bam*HI and *Sac*I by In-Fusion HD Cloning Kit (Takara Bio, Shiga, Japan). For *awf2*, *awf5,* and *awf16*, respectively, a 5.2 kb fragment containing *THO5,* a 4.4 kb fragment containing *CDK8* and a 6.8 kb fragment containing *HSI2/VAL1* were amplified and inserted into pBI101 with *Sal*I and *Sac*I. For *awf4*, *awf9*, and *awf18*, a 5.1 kb fragment containing *FLS2* was amplified and inserted into pBI101 with *Sal*I and *Sac*I.

### Plant transformation

*Agrobacterium tumefaciens* floral dip was used to transform plants, with minor modifications (Clough and Bent 1998). We used *A. tumefacians* strains LBA4404 or GV3101::pMP90 (Koncz and Schell 1986; Weigel and Glazebrook 2002). For constructs in pBIB-HYG, hygromycin B resistant (Hyg^R^) T_1_ plants were selected and grown using standard techniques. We selected T_2_ plants that showed a 3:1 segregation ratio for both hygromycin B resistance and bioluminescence, suggesting they contained a single T-DNA, and obtained T_3_ plants from these selected plants. These T_3_ plants were selected as homozygous by lack of segregation for hygromycin B resistance and bioluminescence. Bulk T_4_ seeds were generated from the selected T_3_ plants. For transformation with pBI101-derived vectors, kanamycin was used for selection.

### Measurement of bioluminescence response

The bioluminescence response of seedlings treated with flg22 was performed as follows. Surface sterilized seeds were incubated at 4°C in the dark for 2 days and sown into wells of a 96-well microplate (Luminunc^TM^ Plates White F96; Thermo Fisher Scientific K.K., Tokyo, Japan) containing 150 μL half-strength liquid MS medium containing 0.5% sucrose and 50 μM D-luciferin-K (Biosynth, Naperville, IL, USA or Promega Japan, Tokyo, Japan) and germinated in continuous light. After 7 days, seedlings were treated with 0.5 μM flg22 (GenScript, Piscataway, NJ, USA), 0.5 μM elf18 (Life Technologies Japan Ltd. Tokyo, Japan), or 1 mg/ml chitin (NA-COS-Y, Yaizu Suisan Kagaku Industry Co., Ltd., Shizuoka, Japan) and 96-well plates were sealed with a plate seal (Excel Scientific, Victorville, CA, USA or Plate Seal T, Sanplatec Corp., Osaka, Japan) instead of plastic cover.

Bioluminescence from each well was measured automatically using a commercially available automated bioluminescence monitoring system (model CL96-4; Churitsu Electric Corp., Nagoya, Aichi, Japan) with a robotic plate conveyor (model CI-08L; Churitsu Electric Corp.). Bioluminescence data was analyzed using commercially available software (SL00-01; Churitsu Electric Corp.).

### Mutagenesis of W29-1-4 reporter line and mutant screening

About 8,000 T_4_ seeds from the W29-1-4 reporter line were mutagenized by treatment with 0.3% (v/v) ethylmethanesulfonate (Merck KGaA, Frankfurter, Darmstadt, Germany) for 15 h at 25°C. M_2_ seeds were collected and grouped into 6 pools, each of which contained seeds from about 1,300 M_1_ plants. About 4,000 seedlings from each M_2_ pool were screened for an altered bioluminescence response following flg22 treatment.

Whole genome sequencing of nine *awf* mutant lines (Supplementary Table S8) showed that Our EMS mutagenesis resulted in 1,000-1,500 nucleotide substitutions incorporated per line. If we extrapolate 1,000 SNPs per line to the total 24,000 lines screened, we expect 24,000,000 SNPs in the 24,000 lines. In the arbitrary 1 kb region of the *Arabidopsis* genome (135 Mb), we expect ∼178 SNPs on average after screening all the lines [178 = 24,000,000/ (135 Mb/1 kb)]. This figure suggests that the majority of the genes of *Arabidopsis* genome has been mutated by our condition of EMS mutagenesis.

In M_2_ screening, we assayed the bioluminescence response from seedlings treated with 0.5 μM flg22 using a high-throughput real-time bioluminescence monitoring system (Churitsu Electric Corp.; http://www.churitsu.co.jp/products/bio/highthroughput.html) equipped with EMCCD (Electron Multiplying Charge-Coupled Device) image sensors (Andor Japan, Tokyo, Japan) essentially as described by Kondo and Ishiura. (1994).

### Generation of F_2_ progeny and whole-genome sequencing

To generate the F_2_ progeny used for bulk sequencing, each mutant was crossed to the W29-1-4 reporter line (the parental line of the mutants) and the resulting F_1_ progeny were self-pollinated to produce F_2_ seeds. Bulk DNA for MutMap analysis was prepared from equal amounts of 30 F_2_ mutant individuals. For whole-genome sequencing, DNA samples were extracted from young leaves with the DNeasy Plant Mini Kit (QIAGEN K.K., Tokyo, Japan). We prepared 11 sequence libraries for WGS; (1), the *WRKY29* reporter line T3_W29-1-4 and *awf3* (read length; 75 bp, paired-end sequencing (PE); (2) *awf1*, *awf2*, *awf5*, *awf8*, *awf9*, *awf14*, *awf16*, *awf19*, and *awf21* mutant lines (150 bp, PE). Sequence libraries for PE short reads were constructed using an Illumina TruSeq DNA LT Sample Prep Kit (Illumina K.K., Tokyo, Japan). The libraries were sequenced on a HiSeq high output (150 bp, PE), NextSeq500 (75 bp, PE), or MiSeq (75 bp, PE). Whole genome sequencing data has been deposited with DDBJ BioProject under PRJDB6767. Whole genome sequencing of the 10 independent mutants indicated that each line had 1,470 ± 266 (mean ± s.d.; range 944-1,795) SNPs relative to the W29-1-4, the wild type reporter strain (Supplementary Table S8).

### MutMap analysis

The W29-1-4 reference sequence was constructed by replacing nucleotides in Col-0 with those of W29-1-4 at the 581 SNP positions identified by aligning 3.4 Gb of Illumina W29-1-4 short reads to the Col-0 reference genome (ftp://ftp.ensemblgenomes.org/pub/release-36/plants/fasta/arabidopsis_thaliana/dna/Arabidopsis_thaliana.TAIR10.dna.toplevel.fa) as previously described (Takagi et al. 2015).

Illumina short reads generated from the bulked DNA samples were filtered by Phred quality score and aligned to the W29-1-4 reference sequence using BWA (Li and Durbin 2009). Alignments were converted to SAM/BAM files using SAMtools (Li et al. 2009), and low-quality SNPs were excluded with a Coval filter (Kosugi et al. 2013). Additionally, SNPs that were detected by self-alignment of the parental W29-1-4 short reads to the W29-1-4 reference sequence were judged to be spurious and were excluded from the analysis. SNP-index was calculated at all SNP positions with Coval (Kosugi et al. 2013). All steps were automatically processed using the MutMap_v1.4.5 pipeline (http://genome-e.ibrc.or.jp/home/bioinformatics-team/mutmap). The mutation responsible to the given phenotype should have SNP-index = 1. However, sometimes NGS short reads contain sequence errors and they may cause SNP-index < 1. Also, there is a possibility that wild-type F_2_ plants were erroneously included in the mutant bulk, which may cause SNP-index < 1. Therefore, we sought the causative SNPs from the SNPs showing SNP-index being close to 1.

#### RNA extraction and qRT-PCR

Gene expression assays were performed on eight day old seedlings and five week old mature leaves. Seedlings and leaves were treated with 0.5 μM flg22 for 60 or 180 min and frozen in liquid nitrogen. Total RNA was extracted using RNeasy Plant Mini kit (QIAGEN K.K.) according to the manufacturer’s instructions. The RNA samples were treated with TURBO DNase (Thermo Fisher Scientific K.K.). cDNA was synthesized using ReverTra Ace® (Toyobo, Osaka, Japan). The qRT-PCR was performed as previously described by Kobayashi et al. (2017). *EF1α* gene (*At1g07920*) was used as internal control. Primers used for quantitative PCR for *EF1α* (Jung et al. 2009), *WRKY29* (Hsu et al. 2013), *FRK1* (He et al. 2006), *CYP81F2* (*At5g57220*), *Luciferase* are listed in Table S7.

#### Bacterial inoculation assay

Inoculation was performed as previously described by Jelenska et al. (2010), Yamada et al. (2016), and Zhang et al. (2017) with some modifications.

Briefly, *Pseudomonas syringae* pv. tomato (*Pto*) *DC3000* was pre-incubated in King’s B (KB) liquid medium with 50 μg/mL rifampicin at 28°C overnight and incubated 4 to 6 hours at 28°C. Freshly grown bacteria from KB agar medium with 50 μg/mL rifampicin. After washing with 10 mM MgSO_4,_ the inoculum was resuspended in 10 mM MgSO_4_ containing 0.01% (v/v) Silwet L-77 to a final OD_600_ = 0.01 and subsequently sprayed on five week old *Arabidopsis*. Inoculated plants were kept in a box covered with plastic for 3 days. Two leaf discs from four or five independent plants were pooled and serially diluted with 10 mM MgSO_4_. Each dilution was plated on LB agar medium containing antibiotics and incubated at 28°C for 2 days.

### Accession numbers

The accession numbers for *Arabidopsis* genes discussed in this article are: *ERF019* (*AT1G22810*), *HSI2/VAL1* (*AT2G30470*), *WRKY29* (*AT4G23550*), *THO5* (*AT5G42920*), *FLS2* (*AT5G46330*), *CDK8* (*AT5G63610*).

## Acknowledgments

We thank Kazue Ito, Aiko Uemura, Eiko Kanzaki, Emiko Sato, and Hideko Kikuchi (IBRC) for technical assistance. Computational analysis was partially performed on the NIG supercomputer at the ROIS National Institute of Genetics.

## Supplemental Material

**Supplementary Fig. S1** *WRKY29* reporter line.

**Supplementary Fig. S2** Bioluminescence patterns of the W29-1-4 reporter line treated with flg22, elf18, or chitin.

**Supplementary Fig. S3** Bioluminescence phenotypes of 22 *awf* mutants as presented in Table 1.

**Supplementary Fig. S4** Box and whisker plot of bioluminescence phenotypes of *awf* mutants as presented in Fig. 3.

**Supplementary Fig. S5** Genetic complementation of *fls2* mutants.

**Supplementary Fig. S6** Segregation of flg22-induced bioluminescence phenotypes in F_2_ progeny.

**Supplementary Fig. S7** MutMap SNP-index plots of nine *awf* mutants.

**Supplementary Fig. S8** Bioluminescence of genetic complementation lines presented in Fig. 6.

**Supplementary Fig. S9** Gene expression analysis in seedlings of *awf* mutants treated with flg22.

**Supplementary Fig. S10** flg22-inducing early ROS production in *awf* mutants.

**Supplementary Fig. S11** MAPK activation assay in *awf* mutants.

**Supplementary Fig. S12** *WRKY29* expression analysis in *awf* mutant leaves treated with flg22.

**Supplementary Table S1** Promoter regions for nine Arabibopsis *Luciferase* reporter lines.

**Supplementary Table S2** Comparison of SNPs between G1042E *fls2* mutants (*awf3*, *awf7,* and *awf9*).

**Supplementary Table S3** Segregation of wild-type (WT) vs. mutant phenotype among the F_2_ progeny of nine *awf* mutants as studied by MutMap.

**Supplementary Table S4** Summary of next generation sequencing data of W29-1-4 and 10 *awf* mutants.

**Supplementary Table S5** Summary of causal regions and number of candidate genes and mutations by MutMap analysis.

**Supplementary Table S6** Candidate causal mutations identified by MutMap.

**Supplementary Table S7** Primers used in this study.

**Supplementary Table S8** Number of reliable SNPs detected between wild type W29-1-4 and 10 *awf* mutant lines.

Supplementary Methods

