## Supplemental Table and Methods for "Lumi-Map, a real-time luciferase bioluminescence screen of mutants combined with MutMap, reveals *Arabidopsis* genes involved in PAMP-triggered immunity"

Table S1.

Promoter regions for nine Arabidopsis *Luciferase* reporter lines.

| **Reporter line** | **Accession No.** | **Promoter region** |
|---|---|---|
| p*WRKY29-LUC* | At4g23550 | -1931 to -1 |
| p*WRKY18-LUC* | At4g31800 | -4270 to -1 |
| p*WRKY28-LUC* | At4g18170 | -3770 to -1 |
| p*RBOHD-LUC* | At5g47910 | -2449 to -1 |
| p*PAL1-LUC* | At2g37040 | -4056 to -1 |
| p*PR4-LUC* | At3g04720 | -3469 to -1 |
| p*ERF1-LUC* | At3g23240 | -4026 to -1 |
| p*At1g51890-LUC* | At1g51890 | -3500 to -1 |
| p*At2g17740-LUC* | At2g17740 | -3000 to -1 |

**Table S2**

**Comparison of SNPs between G1042E *fls2* mutants (*awf3*, *awf7*, and *awf9*)**

| SNPs position | Accession No. | W29-1-4 | *awf3* | *awf7* | *awf9* |
|---|---|---|---|---|---|
| Chr5_18794926 | At5g46330 (*FLS2*) | G | A | A | A |
| Chr1_30359196 | At1g80780 | G | A | A | G |
| Chr2_221717 | At2g01490 | C | T | T | C |
| Chr2_19588296 | At2g47820 | C | T | T | C |
| Chr3_112783 | At3g01320 | G | A/G* | A | G |
| Chr3_23431485 | At3g63460 | G | A | A | G |
| Chr4_136927 | - | G | A/G* | G | G |
| Chr4_12596814 | At4g24290 | C | T | C | C |
| Chr5_9413283 | - | G | A | A | G |
| Chr5_26849650 | At5g67290 | C | T | C | C |
| Chr1_30313605 | At1g80640 | G | G | G | A |
| Chr2_389457 | At2g01860 | C | C | C | T |
| Chr2_19543922 | At2g47670 | C | C | C | T |
| Chr3_3751 | At3g01015 | C | C | C | T |
| Chr3_23445048 | At3g09985 | G | G | G | A |
| Chr4_274913 | At4g00660 | G | G | G | A |
| Chr4_18410304 | At4g39670 | C | C | C | T |
| Chr5_26939360 | At5g67520 | G | G | G | A |

*Both nucleotides were detected by sanger sequencing (hetero).

Green and red letters indicate SNPs specifically detected in *awf3* and *awf9*, respectively.

Table S3

Segregation of wild-type (WT) vs. mutant phenotype among the $F_2$ progeny of nine *awf* mutants as studied by MutMap.

| **Mutant line** | **Phenotype** | **Total number of $F_2$ seedlings** | **WT type** | **Mutant type** | **3:1 segregation $\chi^2$** |
|---|---|---|---|---|---|
| *awf1* | low | 750 | 553 | 197 | 0.64 (n.s.) |
| *awf2* | low | 685 | 538 | 147 | 4.58 ($P < 0.05$) |
| *awf5* | low | 283 | 209 | 74 | 0.20 (n.s.) |
| *awf8* | low | 350 | 265 | 85 | 0.10 (n.s.) |
| *awf9* | low | 703 | 546 | 157 | 2.67 (n.s.) |
| *awf14* | low | 334 | 291 | 43 | 26.19 ($P < 0.01$) |
| *awf16* | low | 352 | 282 | 70 | 4.91 ($P < 0.05$) |
| *awf19* | high | 356 | 293 | 63 | 10.12 ($P < 0.01$) |
| *awf21* | high | 582 | 522 | 60 | 71.77 ($P < 0.01$) |

Table S4

Summary of next generation sequencing of W29-1-4 and ten *awf* mutant lines.

| **Name** | **No. of bulked plant** | **No. of PE reads** | **Read size (bp)** | **Total size (Gbp)** | **Coverage (%)** | **Read depth** | **Purpose** |
|---|---|---|---|---|---|---|---|
| T3_W29-1-4* | - | 22,404,355 | 75 | 3.4 | 96.0 | 18.7 | To make consensus genome |
| *awf1*** | 30 | 25,398,118 | 151 | 7.7 | 97.7 | 35.2 | To perform MutMap analysis |
| *awf2*** | 30 | 30,035,642 | 151 | 9.1 | 97.7 | 45.1 | To perform MutMap analysis |
| *awf5*** | 30 | 8,769,364 | 151 | 2.7 | 97.5 | 13.8 | To perform MutMap analysis |
| *aw8*** | 30 | 10,287,589 | 151 | 3.1 | 97.5 | 16.3 | To perform MutMap analysis |
| *awf9*** | 30 | 27,369,097 | 151 | 8.3 | 97.7 | 41.0 | To perform MutMap analysis |
| *awf14*** | 30 | 9,660,863 | 151 | 2.9 | 97.5 | 15.4 | To perform MutMap analysis |
| *awf16*** | 30 | 11,880,004 | 151 | 3.6 | 97.6 | 19.3 | To perform MutMap analysis |
| *awf19*** | 30 | 15,083,219 | 151 | 4.6 | 97.6 | 23.6 | To perform MutMap analysis |
| *awf21*** | 30 | 10,829,817 | 151 | 3.2 | 97.5 | 16.5 | To perform MutMap analysis |
| *awf3**** | - | 13,380,122 | 75 | 1.9 | 95.4 | 10.1 | To make SNP marker |

* Sequenced by NextSeq 500 platform

** Sequenced by HiSeq highoutput

*** Sequenced by MiSeq platform

Table S5

Summary of causal regions and number of candidate genes and mutations by MutMap analysis.

| **Mutant lines** | **No. of bulked plants** | **Regions of SNP cluster with homozygous SNPs (SNP index ≧0.95)** | **Causal reagions (Mbp)** | **Number of candidate genes** | **Number of intronic mutations** | **Number of mutations upstream of transcription start (in 2 kb)** |
|---|---|---|---|---|---|---|
| *awf1* | 30 | Chr1: 7271229..9026384 | 1.76 | 6 | 1 | 5 |
| *awf2* | 30 | Chr5: 16069066..17741064 | 1.67 | 5 | 2 | 4 |
| *awf5* | 30 | *Chr5: 22435937..25580600 | 3.14 | 5 | 3 | 5 |
| *awf8* | 30 | Chr2: 7369615..11247682 | 3.88 | 18 | 7 | 7 |
| *awf9* | 30 | Chr5: 18794926..19770835 | 0.98 | 3 | 0 | 1 |
| *awf14* | 30 | Chr5: 18795333..22743992 | 3.95 | 17 | 5 | 5 |
| *awf16* | 30 | Chr2: 10418588..14599445 | 4.18 | 9 | 7 | 10 |
| *awf19* | 30 | Chr5: 1322851..6517428 | 5.19 | 20 | 8 | 10 |
| *awf21* | 30 | *Chr1: 10187374..11346652 | 1.16 | 2 | 1 | 1 |

*Region of SNP cluster with SNP index≧0.9

Table S6 Candidate of causal mutations identified by MutMap

| Mutant lines | Chromosome coordinate | Reference base | Altered base | Depth | SNP-index | Accession | Note | Type of mutation |
|---|---|---|---|---|---|---|---|---|
| *awf1* | chr1: 7271229 | C | T | 31 | 0.96 | AT1G20900 | AT-hook motif nuclear-localized protein 27 | -1298 |
| | chr1: 7437722 | C | T | 42 | 0.95 | AT1G21250 | cell wall-associated kinase 1 | -1450 |
| | chr1: 7641919 | C | T | 36 | 1 | AT1G21740 | hypothetical protein | H114Y |
| | chr1: 8074904 | C | T | 29 | 0.96 | AT1G22810 | ERF019 | W22* |
| | chr1: 8163791 | C | T | 52 | 1 | AT1G23040 | hydroxyproline-rich glycoprotein family protein | -992 |
| | chr1: 8223678 | C | T | 38 | 0.97 | AT1G23190 | phosphoglucomutase 3 | A292V |
| | chr1: 8468863 | C | T | 32 | 0.96 | AT1G23940 | apoptosis inhibitory protein | intron |
| | chr1: 8553760 | C | T | 39 | 1 | AT1G24160 | uncharacterized protein | R511Q |
| | chr1: 8645637 | C | T | 29 | 1 | AT1G24380 | no-apical-meristem-associated carboxy-terminal domain protein | -420 |
| | chr1: 8673408 | C | T | 48 | 1 | AT1G24470 | beta-ketoacyl reductase 2 | -507 |
| | chr1: 8755132 | C | T | 23 | 1 | AT1G24706 | AtTHO2 | A1785V |
| | chr1: 9026384 | C | T | 30 | 1 | AT1G26110 | DCP5 | A119T |
| *awf2* | chr5: 16167341 | C | T | 43 | 0.95 | AT5G40400 | pentatricopeptide repeat-containing protein | R300C |
| | chr5: 16357820 | C | T | 45 | 0.95 | AT5G40840 | SYN2 | -1512 |
| | chr5: 16373616 | C | T | 62 | 0.98 | AT5G40870 | uridine kinase | -1184 |
| | chr5: 16438686 | C | T | 41 | 0.95 | AT5G41070 | DRB5 | P47L |
| | chr5: 16811152 | C | T | 45 | 0.95 | AT5G42020 | luminal binding protein | -505 |
| | chr5: 16955490 | C | T | 64 | 1 | AT5G42400 | ATXR7 | intron |
| | chr5: 17033201 | G | A | 46 | 0.97 | AT5G42590 | CYP71A16 | intron |
| | chr5: 17207377 | C | T | 47 | 1 | AT5G42920 | AtTHO5 | W405* |
| | chr5: 17538254 | G | A | 44 | 0.97 | AT5G43660 | 2-oxoglutarate (2OG) and Fe(II)-dependent oxygenase superfamily | P39S |
| | chr5: 17589113 | G | A | 41 | 0.97 | AT5G43770 | proline-rich family protein | E103K |
| | chr5: 17741064 | G | A | 51 | 0.96 | AT5G44080 | bZIP transcription factor family protein | -1114 |
| *awf5* | chr5: 22435937 | C | T | 16 | 1 | AT5G55320 | putative long-chain-alcohol O-fatty-acyltransferase 7 | G255R |
| | chr5: 22514001 | C | T | 12 | 1 | AT5G55565 | defensin-like protein | -412 |
| | chr5: 23023555 | C | T | 15 | 0.93 | AT5G56900 | CwfJ-like family protein | -789 |
| | chr5: 23683831 | C | T | 16 | 1 | AT5G58600 | Pmr5/Cas1p | intron |
| | chr5: 24143314 | C | T | 8 | 1 | AT5G59960 | uncharacterized protein | V206M |
| | chr5: 24172720 | C | T | 12 | 1 | AT5G60030 | hypothetical protein | D81N |
| | chr5: 24173172 | C | T | 13 | 1 | AT5G60030 | hypothetical protein | -125 |
| | chr5: 24210269 | C | T | 19 | 0.94 | AT5G60120 | target of early activation tagged 2 | intron |
| | chr5: 24353414 | C | T | 20 | 0.95 | AT5G60580 | RING/U-box superfamily protein | -262 |
| | chr5: 24769820 | C | T | 23 | 0.95 | AT5G61610 | Oleosin family protein | -227 |
| | chr5: 25308787 | C | T | 13 | 0.92 | AT5G63090 | LATERAL ORGAN BOUNDARIES | G166E |
| | chr5: 25464254 | C | T | 9 | 1 | AT5G63610 | cyclin-dependent kinase | W268* |
| | chr5: 25580600 | C | T | 13 | 1 | AT5G63920 | topoisomerase 3alpha | intron |
| *awf8* | chr2: 7369615 | G | A | 28 | 0.96 | AT2G16970 | MEE15 | E23K |
| | chr2: 7417716 | G | A | 22 | 1 | AT2G17055 | Toll-Interleukin-Resistance domain family protein | -936 |
| | chr2: 7425296 | G | A | 16 | 1 | AT2G17060 | Disease resistance protein | M724I |
| | chr2: 7444004 | G | A | 15 | 1 | AT2G17110 | DNA-directed RNA polymerase subunit beta | P522S |
| | chr2: 7955072 | G | A | 28 | 1 | AT2G18300 | HOMOLOG OF BEE2 INTERACTING WITH IBH 1 | -286 |
| | chr2: 8040614 | G | A | 9 | 1 | AT2G18530 | Protein kinase superfamily protein | G159D |
| | chr2: 8176427 | G | A | 16 | 1 | AT2G18880 | VIN3-like 2 | intron |
| | chr2: 8326715 | G | A | 13 | 1 | AT2G19190 | FRK1 | P692S |
| | chr2: 8465961 | G | A | 19 | 1 | AT2G19560 | ENHANCED ETHYLENE RESPONSE 5 | P402L |
| | chr2: 8637368 | G | A | 15 | 1 | AT2G20000 | CDC27b | -67 |
| | chr2: 8738711 | G | A | 19 | 1 | AT2G20270 | Thioredoxin superfamily protein | intron |
| | chr2: 8789817 | G | A | 19 | 1 | AT2G20362 | transmembrane protein | G16R |
| | chr2: 8799849 | G | A | 14 | 1 | AT2G20400 | myb-like HTH transcriptional regulator family | E76K |
| | chr2: 8807034 | G | A | 19 | 1 | AT2G20420 | succinyl-CoA ligase beta subunit | A236T |
| | chr2: 8832569 | G | A | 17 | 1 | AT2G20490 | NOP10 | intron |
| | chr2: 9045699 | G | A | 20 | 1 | AT2G21090 | PPR-like superfamily protein | T597I |
| | chr2: 9171462 | G | A | 16 | 1 | AT2G21420 | IBR domain containing protein | V375I |
| | chr2: 9202919 | G | A | 9 | 1 | AT2G21480 | Malectin/receptor-like protein kinase family | P817L |
| | chr2: 9215038 | G | A | 24 | 1 | AT2G21520 | Sec14p-like phosphatidylinositol transfer family protein | -482 |
| | chr2: 9532224 | G | A | 25 | 1 | AT2G22450 | RIBA2 | G349E |
| | chr2: 9850420 | G | A | 12 | 1 | AT2G23142 | Plant self-incompatibility protein S1 family | G82E |
| | chr2: 9906629 | G | A | 21 | 0.95 | AT2G23290 | MYB70 | -814 |
| | chr2: 10303085 | C | T | 16 | 1 | AT2G24230 | Leucine-rich repeat protein kinase family | G486S |

| | | | | | | | |
|---|---|---|---|---|---|---|---|
| | chr2: 10320096 | G | A | 9 | 1 | AT2G24260 | LRL1 | intron |
| | chr2: 10340297 | G | A | 21 | 1 | AT2G24300 | Calmodulin-binding protein | -489 |
| | chr2: 10633434 | G | A | 18 | 1 | AT2G25010 | Aminotransferase-like | A473T |
| | chr2: 10722838 | G | A | 12 | 1 | AT2G25170 | PKL | G1156E |
| | chr2: 10778102 | G | A | 22 | 1 | AT2G25310 | ER membrane protein complex subunit-like protein | intron |
| | chr2: 10959026 | G | A | 20 | 1 | AT2G25730 | zinc finger FYVE domain protein | intron |
| | chr2: 11236205 | G | A | 14 | 1 | AT2G26410 | IQ-domain 4 | L134F |
| | chr2: 11703710 | G | A | 11 | 1 | AT2G27350 | OTU-like cysteine protease family protein | intron |
| | chr2: 11780552 | G | A | 18 | 1 | AT2G27600 | SKD1 | -345 |
| *awf9* | chr5: 18794926 | G | A | 32 | 1 | AT5G46330 | FLS2 | G1042E |
| | chr5: 19242248 | G | A | 44 | 1 | AT5G47435 | formyltetrahydrofolate deformylase | G111S |
| | chr5: 19748482 | G | A | 48 | 0.95 | AT5G48690 | Ubiquitin-like superfamily protein | P73L |
| | chr5: 19770835 | G | A | 48 | 1 | AT5G48740 | Leucine-rich repeat protein kinase family protein | -1465 |
| *awf14* | chr5: 18795333 | C | T | 23 | 1 | AT5G46070 | FLS2 | L1150F |
| | chr5: 18798902 | C | T | 14 | 1 | AT5G46340 | REDUCED WALL ACETYLATION 1 | intron |
| | chr5: 18812961 | C | T | 19 | 0.95 | AT5G46370 | ATKCO2 | -1090 |
| | chr5: 18830474 | C | T | 18 | 1 | AT5G46420 | 16S rRNA processing protein RimM family | P100S |
| | chr5: 18933025 | C | T | 16 | 1 | AT5G46660 | protein kinase C-like zinc finger protein | S282F |
| | chr5: 19281981 | C | T | 13 | 1 | AT5G47530 | Auxin-responsive family protein | P171S |
| | chr5: 19398879 | C | T | 19 | 1 | AT5G47910 | RBOHD | intron |
| | chr5: 19610029 | C | T | 9 | 1 | AT5G48385 | FRIGIDA-like protein | L187F |
| | chr5: 19649194 | C | T | 16 | 1 | AT5G48490 | Bifunctional inhibitor/lipid-transfer protein | -831 |
| | chr5: 19656268 | C | T | 13 | 1 | AT5G48510 | BTB/POZ domain-containing protein | -1813 |
| | chr5: 19794162 | C | T | 8 | 1 | AT5G48820 | ICK6 | R62H |
| | chr5: 19921618 | C | T | 13 | 1 | AT5G49140 | Disease resistance protein family | intron |
| | chr5: 20042406 | C | T | 9 | 1 | AT5G49430 | WD40/YVTN repeat and Bromo-WDR9-I-like domain-containing p | junction |
| | chr5: 20057564 | C | T | 8 | 1 | AT5G49460 | ATP citrate lyase subunit B 2 | A455V |
| | chr5: 20532385 | G | A | 13 | 1 | AT5G50420 | O-fucosyltransferase family protein | T84I |
| | chr5: 20778320 | G | A | 18 | 1 | AT5G51110 | Transcriptional coactivator/pterin dehydratase | A220V |
| | chr5: 20855680 | G | A | 18 | 1 | AT5G51310 | 2-oxoglutarate and Fe(II)-dependent oxygenase superfamily protein | -867 |
| | chr5: 21115446 | G | A | 12 | 1 | AT5G51980 | Transducin/WD40 repeat-like superfamily protein | L153F |
| | chr5: 21483308 | G | A | 14 | 1 | AT5G52980 | ER-based factor for assembly of V-ATPase | intron |
| | chr5: 21499198 | G | A | 20 | 0.95 | AT5G53020 | Ribonuclease P protein subunit P38-like protein | intron |
| | chr5: 21590807 | G | A | 19 | 1 | AT5G53210 | SPEECHLESS | -1551 |
| | chr5: 22094263 | C | T | 13 | 1 | AT5G54410 | hypothetical protein (DUF295) | G215D |
| | chr5: 22185756 | C | T | 12 | 1 | AT5G54610 | ankyrin | junction |
| | chr5: 22449297 | C | T | 9 | 1 | AT5G55390 | ENHANCED DOWNY MILDEW 2 | E917K |
| | chr5: 22506734 | C | T | 9 | 1 | AT5G55560 | Protein kinase superfamily protein | E230K |
| | chr5: 22678825 | C | T | 12 | 1 | AT5G56000 | HEAT SHOCK PROTEIN 81.4 | E293K |
| | chr5: 22743992 | C | T | 10 | 1 | AT5G56190 | Transducin/WD40 repeat-like superfamily protein | T225I |
| *awf16* | chr2: 10418588 | G | A | 13 | 1 | AT2G24520 | plasma membrane H+-ATPase | G699R |
| | chr2: 10689156 | G | A | 22 | 0.95 | AT2G25120 | Bromo-adjacent homology (BAH) domain-containing protein | -779 |
| | chr2: 10764824 | G | A | 16 | 1 | AT2G25280 | AmmeMemoRadiSam system protein B | -13 |
| | chr2: 10965533 | G | A | 14 | 0.92 | AT2G25730 | zinc finger FYVE domain protein | T1047I |
| | chr2: 11186583 | G | A | 24 | 1 | AT2G26270 | BRCT domain DNA repair protein | R324H |
| | chr2: 11200301 | G | A | 30 | 0.96 | AT2G26300 | GP ALPHA 1 | intron |
| | chr2: 11244412 | G | A | 18 | 1 | AT2G26430 | arginine-rich cyclin 1 | intron |
| | chr2: 11619696 | G | A | 24 | 0.95 | AT2G27180 | hypothetical protein | -381 |
| | chr2: 11712141 | G | A | 24 | 0.91 | AT2G27380 | extensin proline-rich 1 | -1271 |
| | chr2: 11823259 | G | A | 17 | 1 | AT2G27740 | RAB6-interacting golgin (DUF662) | intron |
| | chr2: 12460255 | G | A | 12 | 1 | AT2G29000 | Leucine-rich repeat protein kinase family protein | -527 |
| | chr2: 12494603 | G | A | 18 | 1 | AT2G29080 | FTSH protease 3 | -1273 |
| | chr2: 12887113 | G | A | 23 | 0.95 | AT2G30200 | EMB3147 | -1336 |
| | chr2: 12983531 | G | A | 17 | 0.94 | AT2G30470 | high-level expression of sugar-inducible gene 2 | Q251* |
| | chr2: 13075055 | G | A | 19 | 1 | AT2G30680 | callose synthase-like protein | intron |
| | chr2: 13096584 | G | A | 23 | 1 | AT2G30740 | Protein kinase superfamily protein | intron |
| | chr2: 13191424 | G | A | 23 | 1 | AT2G30990 | arginine N-methyltransferase, putative (DUF688) | -965 |
| | chr2: 13257306 | G | A | 12 | 1 | AT2G31100 | alpha/beta-Hydrolases superfamily protein | A226V |
| | chr2: 13548684 | G | A | 20 | 1 | AT2G31865 | poly(ADP-ribose) glycohydrolase 2 | P161L |
| | chr2: 13598464 | G | A | 17 | 1 | AT2G31960 | glucan synthase-like 3 | V1525M |
| | chr2: 13633782 | G | A | 15 | 1 | AT2G32030 | Acyl-CoA N-acyltransferases superfamily protein | -230 |
| | chr2: 13887628 | G | A | 17 | 1 | AT2G32740 | galactosyltransferase 13 | R88Q |

| | | | | | | | | |
|---|---|---|---|---|---|---|---|---|
| | chr2: 13939261 | G | A | 15 | 1 | AT2G32850 | Protein kinase superfamily protein | -415 |
| | chr2: 14020745 | G | A | 19 | 1 | AT2G33040 | ATP3 | intron |
| | chr2: 14268949 | G | A | 20 | 0.95 | AT2G33735 | Chaperone DnaJ-domain superfamily protein | intron |
| | chr2: 14441517 | G | A | 21 | 0.95 | AT2G34200 | RING/FYVE/PHD zinc finger superfamily protein | G138D |
| *awf19* | chr5: 1352792 | G | A | 25 | 0.96 | AT5G04690 | Ankyrin repeat family protein | -212 |
| | chr5: 1364831 | G | A | 22 | 0.95 | AT5G04730 | Ankyrin-repeat containing protein | S361F |
| | chr5: 1386620 | G | A | 14 | 1 | AT5G04800 | Ribosomal S17 family protein | -1290 |
| | chr5: 1575648 | G | A | 16 | 1 | AT5G05320 | FAD/NAD(P)-binding oxidoreductase family protein | intron |
| | chr5: 1579278 | G | A | 13 | 1 | AT5G05340 | Peroxidase superfamily protein | S280F |
| | chr5: 1660143 | G | A | 23 | 0.95 | AT5G05570 | transducin family protein / WD-40 repeat family | S368N |
| | chr5: 1829644 | G | A | 14 | 1 | AT5G06070 | RABBIT EARS | -457 |
| | chr5: 1870439 | G | A | 19 | 1 | AT5G06170 | sucrose-proton symporter 9 | D217N |
| | chr5: 2344789 | G | A | 28 | 1 | AT5G07400 | forkhead-associated domain-containing protein | R908K |
| | chr5: 2536624 | G | A | 25 | 1 | AT5G07940 | dentin sialophosphoprotein-like protein | W599* |
| | chr5: 2768070 | G | A | 18 | 1 | AT5G08550 | increased level of polyploidy1-1D | S450F |
| | chr5: 2774827 | G | A | 20 | 1 | AT5G08560 | transducin family protein | -112 |
| | chr5: 2797223 | G | A | 17 | 1 | AT5G08620 | STRESS RESPONSE SUPPRESSOR 2 | intron |
| | chr5: 3011572 | G | A | 22 | 0.95 | AT5G09711 | transmembrane protein | -424 |
| | chr5: 3236132 | G | A | 25 | 1 | AT5G10290 | leucine-rich repeat transmembrane protein kinase | A421V |
| | chr5: 3585968 | G | A | 25 | 0.96 | AT5G11240 | transducin family protein | intron |
| | chr5: 3778512 | G | A | 32 | 0.96 | AT5G11720 | Glycosyl hydrolases family 31 | D454N |
| | chr5: 3788080 | G | A | 28 | 1 | AT5G11750 | Ribosomal protein L19 family protein | intron |
| | chr5: 4005088 | G | A | 28 | 0.92 | AT5G12370 | exocyst complex component sec10 | A578V |
| | chr5: 4143957 | G | A | 14 | 1 | AT5G13060 | ARMADILLO BTB protein 1 | A194T |
| | chr5: 4298019 | G | A | 22 | 1 | AT5G13400 | peptide transporter-like protein | A266V |
| | chr5: 4359988 | G | A | 27 | 1 | AT5G13550 | sulfate transporter 4.1 | -295 |
| | chr5: 4390419 | G | A | 12 | 1 | AT5G13630 | ABA-BINDING PROTEIN | intron |
| | chr5: 4694188 | G | A | 36 | 1 | AT5G14550 | Core-2/I-branching beta-1,6-N-acetylglucosaminyltransferase famil | -132 |
| | chr5: 4713135 | G | A | 24 | 0.95 | AT5G14610 | DEAD box RNA helicase family protein | V238I |
| | chr5: 5320381 | G | A | 16 | 1 | AT5G16270 | sister chromatid cohesion 1 protein 4 | R575H |
| | chr5: 5469355 | G | A | 32 | 1 | AT5G16680 | RING/FYVE/PHD zinc finger superfamily protein | Q858* |
| | chr5: 5482834 | G | A | 24 | 1 | AT5G16700 | Glycosyl hydrolase superfamily protein | G419E |
| | chr5: 5499504 | G | A | 17 | 1 | AT5G16730 | Encodes a microtubule-associated protein | E431K |
| | chr5: 5603707 | G | A | 18 | 1 | AT5G17030 | UDP-glucosyl transferase 78D3 | P291S |
| | chr5: 5650215 | G | A | 24 | 1 | AT5G17170 | ENH1 | intron |
| | chr5: 5723852 | G | A | 18 | 1 | AT5G17370 | Transducin/WD40 repeat-like superfamily protein | intron |
| | chr5: 5843508 | G | A | 23 | 1 | AT5G17710 | embryo defective 1241 | -1868 |
| | chr5: 5902825 | G | A | 18 | 0.94 | AT5G17860 | calcium exchanger 7 | P509L |
| | chr5: 6090978 | G | A | 17 | 1 | AT5G18390 | Pentatricopeptide repeat (PPR) superfamily | G9S |
| | chr5: 6204327 | G | A | 28 | 0.96 | AT5G18630 | alpha/beta-Hydrolases superfamily protein | intron |
| | chr5: 6359348 | G | A | 22 | 1 | AT5G19030 | RNA-binding family protein | -189 |
| | chr5: 6517428 | G | A | 30 | 1 | AT5G19350 | RNA-binding family protein | -1350 |
| *awf21* | chr1: 10448155 | C | T | 14 | 0.92 | AT1G29850 | double-stranded DNA-binding family protein | intron |
| | chr1: 10511574 | C | T | 11 | 0.9 | AT1G30000 | MNS3 | G84R |
| | chr1: 10595608 | C | T | 18 | 0.94 | AT1G30135 | TIFY5A | -745 |
| | chr1: 10920076 | C | T | 11 | 0.9 | AT1G30760 | FAD-binding Berberine family protein | P414S |

Red cells indicate the mutasions in causal gene of each mutant.

Gray cells indicate the intronic mutations (intron) and mutations in 2-kb upstream of transcription start site (position from the start site).

Table S7 Primers used in this study

| **Primer** | **Sequence (5' to 3')** | **Description** |
|---|---|---|
| AtWRKY29-SacI-F | AAGAGCTCCGAACTCACAGAAGTCAATA | for p*WRKY29* reporter construction |
| AtWRKY29-NcoI-R | AAACCATGGATAAGCCACCTCACCCATAT | for p*WRKY29* reporter construction |
| KpnI-Luc-R | TTGGTACCGGATCCGATCTAGTAACATAG | for reporter construction |
| awf1_AT1G22810_F_BamHI | GGGGGATCCCTGAAACACTATAAACGCGTGCC | for pBI101-ERF019 (AT1G22810) |
| awf1_AT1G22810_R_SacI | ACCGAGCTCTCAAACGTGATCGTGGCCGCCCAG | for pBI101-ERF019 (AT1G22810) |
| awf2_AT5G42920_IF_F | ATGCCTGCAGGTCGACTTCAAAAAGAAGGATGGCCTAAAG | for pBI101-THO5 (AT5G42920) |
| awf2_AT5G42920_IF_R | GATCGGGGAAATTCGCTAGCATGGATAACCCGATGCAC | for pBI101-THO5 (AT5G42920) |
| awf5_AT5G63610_IF_F | ATGCCTGCAGGTCGAATGAAATTCTCATCAAAACCAAAGG | for pBI101-CDK8 (AT5G63610) |
| awf5_AT5G63610_IF_R | GATCGGGGAAATTCGTTAGAGGCGTCTGGATTTGTTAGG | for pBI101-CDK8 (AT5G63610) |
| awf16_AT2G30470_IF_F | ATGCCTGCAGGTCGAAAAAAAATGTCTATTGCAAGCTGG | for pBI101-HSI2/Val1(AT2G30470) |
| awf16_AT2G30470_IF_R | GATCGGGGAAATTCGTCAGCTTGAAACTCTCGGCTCTTC | for pBI101-HSI2/Val1(AT2G30470) |
| FLS2_C_IF_F | ATGCCTGCAGGTCGACGCCAGATTTAGGCTCTGGTCCG | for pBI101-FLS2 |
| FLS2_C_IF_R | GATCGGGGAAATTCGCTAAACTTCTCGATCCTCGTTACG | for pBI101-FLS2 |
| WRKY29_qRT_F1 | CGGAGATGGAGACAAGTGGCTT | for WRKY29 qPCR |
| WRKY29_qRT_R1 | TGTGAGGATCGTTTGTGTGGAGAA | for WRKY29 qPCR |
| Luc_qRT_F | TGAGTACTTCGAAATGTCCGTTC | for Luciferase qPCR |
| Luc_qRT_R | GTATTCAGCCCATATCGTTTCAT | for Luciferase qPCR |
| EF1α_qRT_F | AGGTCCACCAACCTTGACTG | for EF1α qPCR |
| EF1α_qRT_R | GAGACTCGTGGTGCATCTCA | for EF1α qPCR |
| AtWRKY18-F | TTTAAACGAATTCGCGTCGACGAAACGTCGGGTTAATCAGAATTTC | for p*WRKY18* reporter construction |
| AtWRKY18-R | TTTGGCGTCTTCCATAAAAGAAACCTTTATCTTAAGATAC | for p*WRKY18* reporter construction |
| AtWRKY28-F | TTTAAACGAATTCGCGTCGACCCAAAAGTTATCCTCTCCTTTCTC | for p*WRKY28* reporter construction |
| AtWRKY28-F | TTTGGCGTCTTCCATGGTGAAGAACAATGAAGAGAGAGG | for p*WRKY28* reporter construction |
| AtRBOHD-F | TTTAAACGAATTCGCCTGCAGATATAAGCAAAGCCTTTTGTCG | for p*RBOHD* reporter construction |
| AtRBOHD-F | TTTGGCGTCTTCCATCGAATTCGAGAAACCAAAAAGATC | for p*RBOHD* reporter construction |
| AtPAL1-F | TTTAAACGAATTCGCCTGCAGTCCTTAGATATAAGATATCAATC | for p*PAL1* reporter construction |
| AtPAL1-R | TTTGGCGTCTTCCATTTAGACTTTTGATCTTAGTTTTAC | for p*PAL1* reporter construction |
| AtPR4-F | TTTAAACGAATTCGCCTGCAGTTAGGTTGAGAGTCTGTTTGATT | for p*PR4* reporter construction |

| | | |
|---|---|---|
| AtPR4-R | TTTGGCGTCTTCCATGATCGATAAGTCTTTGTTTTCTTG | for p*PR4* reporter construction |
| AtERF1-F | TTTAAACGAATTCGCGTCGACATGGTAACTTTTAACAATTATTAGT | for p*ERF1* reporter construction |
| AtERF1-R | TTTGGCGTCTTCCATGTAGAAAAAATACTCTGTTTCTTGA | for p*ERF1* reporter construction |
| At1g51890-F | TTTAAACGAATTCGCCTGCAGTTAGAAATATTAAGGCCGGAATC | for p*At1g51890* reporter construction |
| At1g51890-R | TTTGGCGTCTTCCATGTTTTTTGGGGATGTTTGTTCTG | for p*At1g51890* reporter construction |
| At2g17740-F | TTTAAACGAATTCGCCTGCAGTCTTACTTAGTTCTCTTATAATT | for p*At2g17740* reporter construction |
| At2g17740-R | TTTGGCGTCTTCCATTAGAGGTAATATTCTTGATTTGC | for p*At2g17740* reporter construction |

Table S8

Number of reliable SNPs detected between wild type W29-1-4 and ten *awf* mutant lines.

“W29-1-4” reference sequence is deposited to DDBJ BioProjcet PRJDB6767.

| **Reporter line** | **Reference sequence** | **Number of SNPs detected** |
|---|---|---|
| W29-1-4 | Col-0* | 581 |

* Arabidopsis_thaliana.TAIR10

Number of reliable SNPs detected between ten *awf* mutant lines.

| **Mutant lines** | **Reference sequence** | **Number of SNPs detected** | **(G to A or C to T transition)** |
|---|---|---|---|
| *awf1* | W29-1-4 | 1540 | (1263) |
| *awf2* | W29-1-4 | 1594 | (1327) |
| *awf3* | W29-1-4 | 1708 | (1357) |
| *awf5* | W29-1-4 | 944 | (760) |
| *awf8* | W29-1-4 | 1766 | (1537) |
| *awf9* | W29-1-4 | 1361 | (1098) |
| *awf14* | W29-1-4 | 1795 | (1572) |
| *awf16* | W29-1-4 | 1451 | (1214) |
| *awf19* | W29-1-4 | 1242 | (981) |
| *awf21* | W29-1-4 | 1301 | (1046) |

**Supplemental Methods**

***ROS assay***

ROS released by leaf tissue was assayed as described (Ranf et al. 2011) using 3 mm leaf discs in 96 well plates containing 0.1 mL distilled water with 200 μM luminal and 10 μg/mL horseradish peroxidase (Sigma-Aldrich Japan K.K. Tokyo, Japan) measured in 2 minute intervals for 60 minutes using Luminoskan Ascent 2.1. (Thermo Scientific K.K.).

***MAPK activation assay***

MAPK activation assays were performed on eight day old seedlings grown in liquid medium. Seedlings were then elicited with 0.5 μM flg22 for 10 or 15 min and frozen in liquid nitrogen. MAPK activation was monitored by western blot with antibodies that recognize the dual phosphorylation of the activation loop of MAPK (pTEpY). Phospho-p44/42 MAPK (Erk1/2; Thr-202/Tyr-204) rabbit monoclonal antibodies were used according to the manufacturer's protocol (#9101, Cell Signaling Technology Japan, K.K. Tokyo, Japan). Blots were stained with Ponceau S to verify equal loading.

**Plasmid construction for *Luciferase* reporters**

The defense-related reporter gene cassettes, $P_{WRKY18}$::$LUC^{+}$, $P_{WRKY28}$::$LUC^{+}$, $P_{RBOHD}$::$LUC^{+}$, $P_{PAL1}$::$LUC^{+}$, $P_{PR4}$::$LUC^{+}$, $P_{ERF1}$::$LUC^{+}$, $P_{At1g51890}$::$LUC^{+}$, or $P_{At2g17740}$::$LUC^{+}$ were inserted into pBIB-HYG (Becker et al. 1990) with *Sal*I and *Sac*I, respectively.
