## Supplemental Figures for "Lumi-Map, a real-time luciferase bioluminescence screen of mutants combined with MutMap, reveals *Arabidopsis* genes involved in PAMP-triggered immunity"

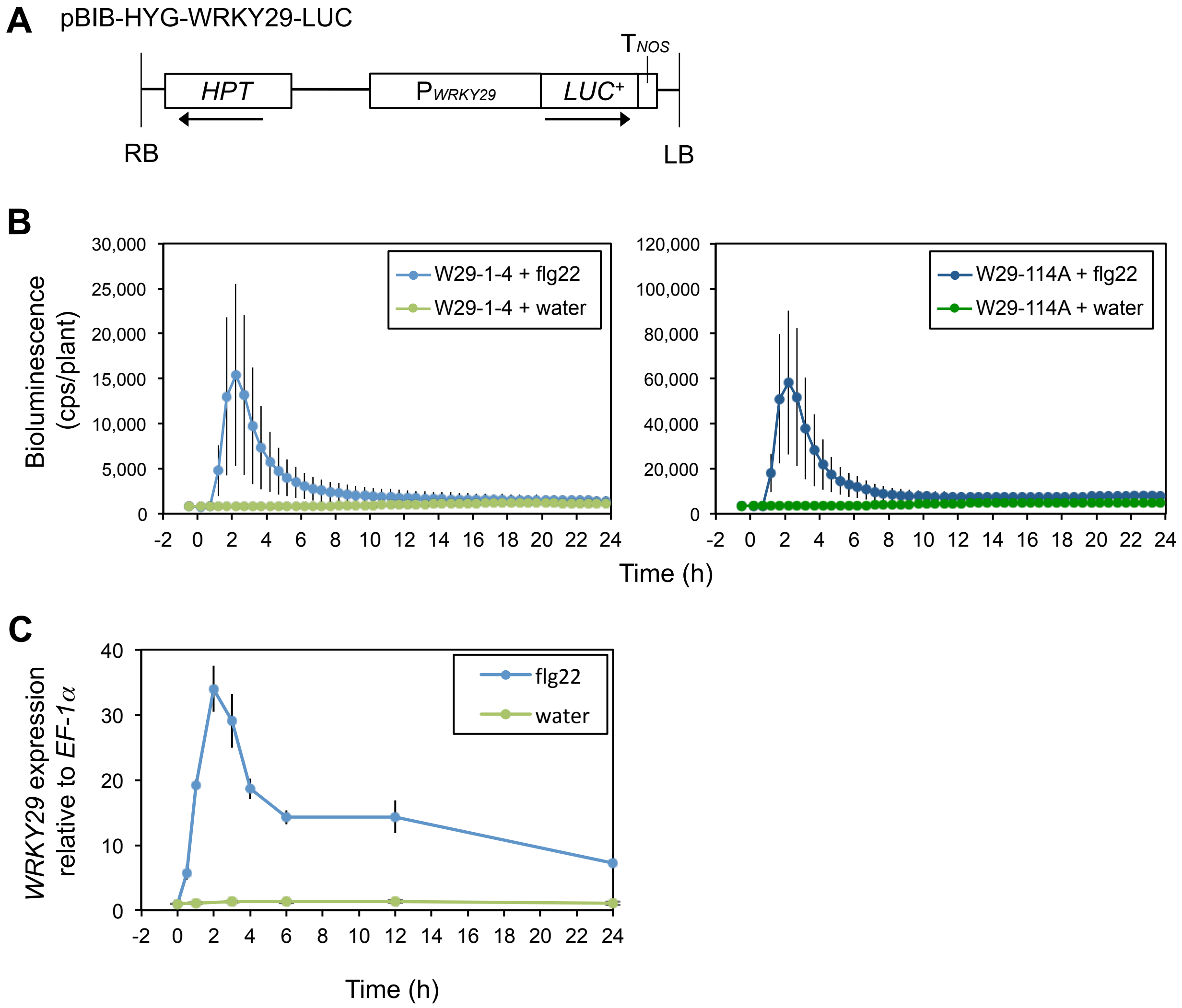


**Fig. S1** *WRKY29* reporter line. **A** Structure of the $P_{WRKY29}$::*LUC*+ reporter construct. *HPT*, a hygromycine resistance gene cassette in pBIB-HYG; $P_{WRKY29}$, promoter of *WRKY29* (-1931 to -1); *LUC*+, coding region of a modified *luciferase* gene derived from *Photinus pyralis*; $T_{NOS}$, transcriptional terminator of *NOS* (*nopaline synthase*) from *Agrobacterium*; LB, left-border sequence of T-DNA; RB, right-border sequence of T-DNA. The transcriptional direction of gene cassettes are shown by arrows. **B** Luciferase-mediated bioluminescence patterns of *WRKY29* reporter lines. Eight day old seedlings of W29-1-4 and W29-114A were treated with water or 0.5 μM flg22. Bioluminescence from each seedling was monitored with a real-time bioluminescence monitoring system at the indicated time points. Data are shown as mean ± SE from at least 13 seedlings per treatment. **C** *WRKY29* expression levels in flg22-treated Arabidopsis seedlings (Col-0) as measured by qRT-PCR. Data are shown as mean ± SE from at least 30 seedlings per treatment.

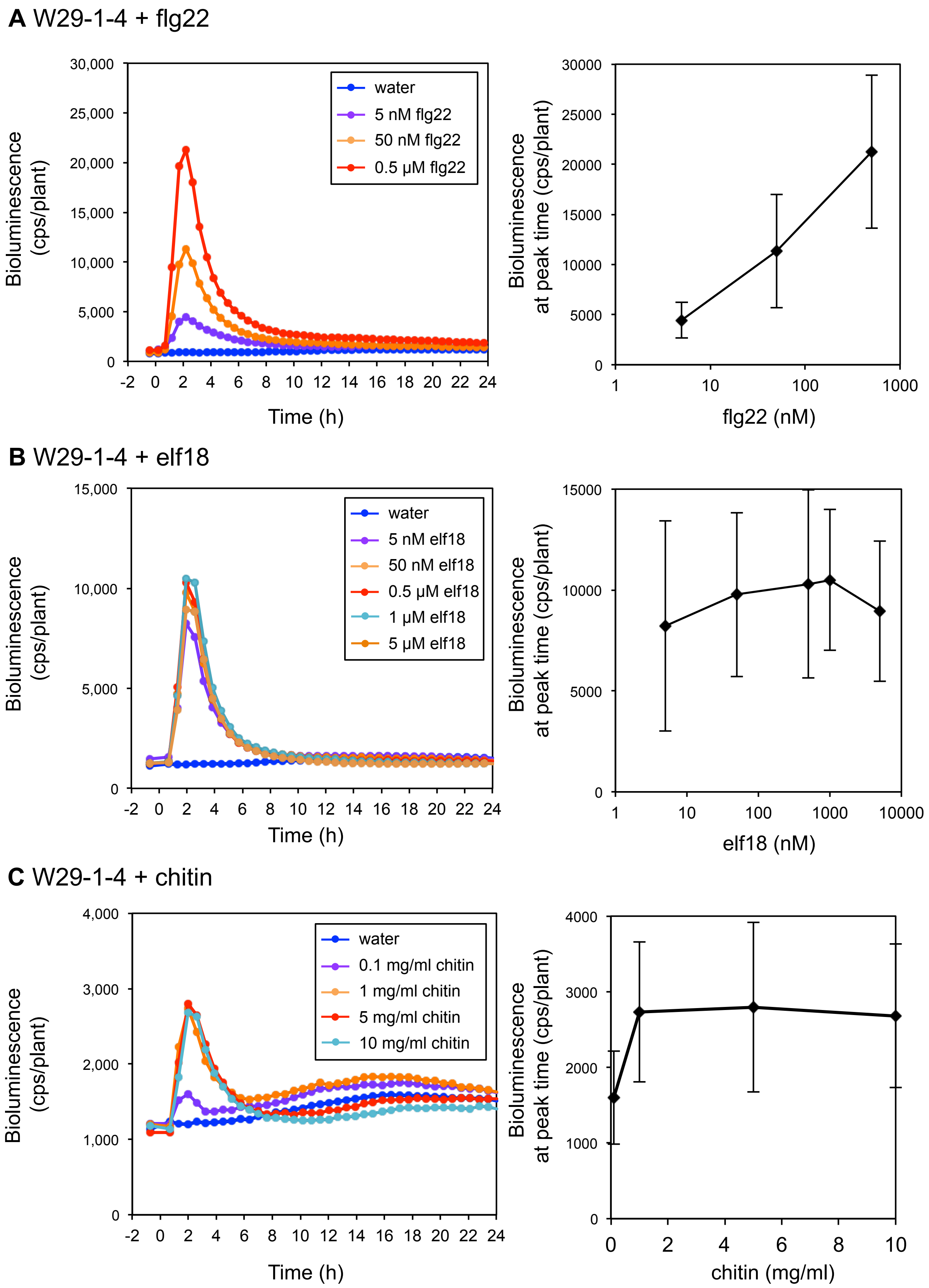


**Fig. S2** Bioluminescence patterns of the W29-1-4 reporter line treated with flg22, elf18, or chitin. Eight day old W29-1-4 seedlings were treated with water, flg22 (**A**), elf18 (**B**), or chitin (**C**) at the indicated concentrations. Bioluminescence from each seedling was monitored at the indicated time points (left panels). Induction strength varies depending on the ligand concentration (right panels). Bioluminescence at the peak time point was compared across the indicated concentration range. Data are shown as mean ± SE from at least 13 seedlings per treatment.

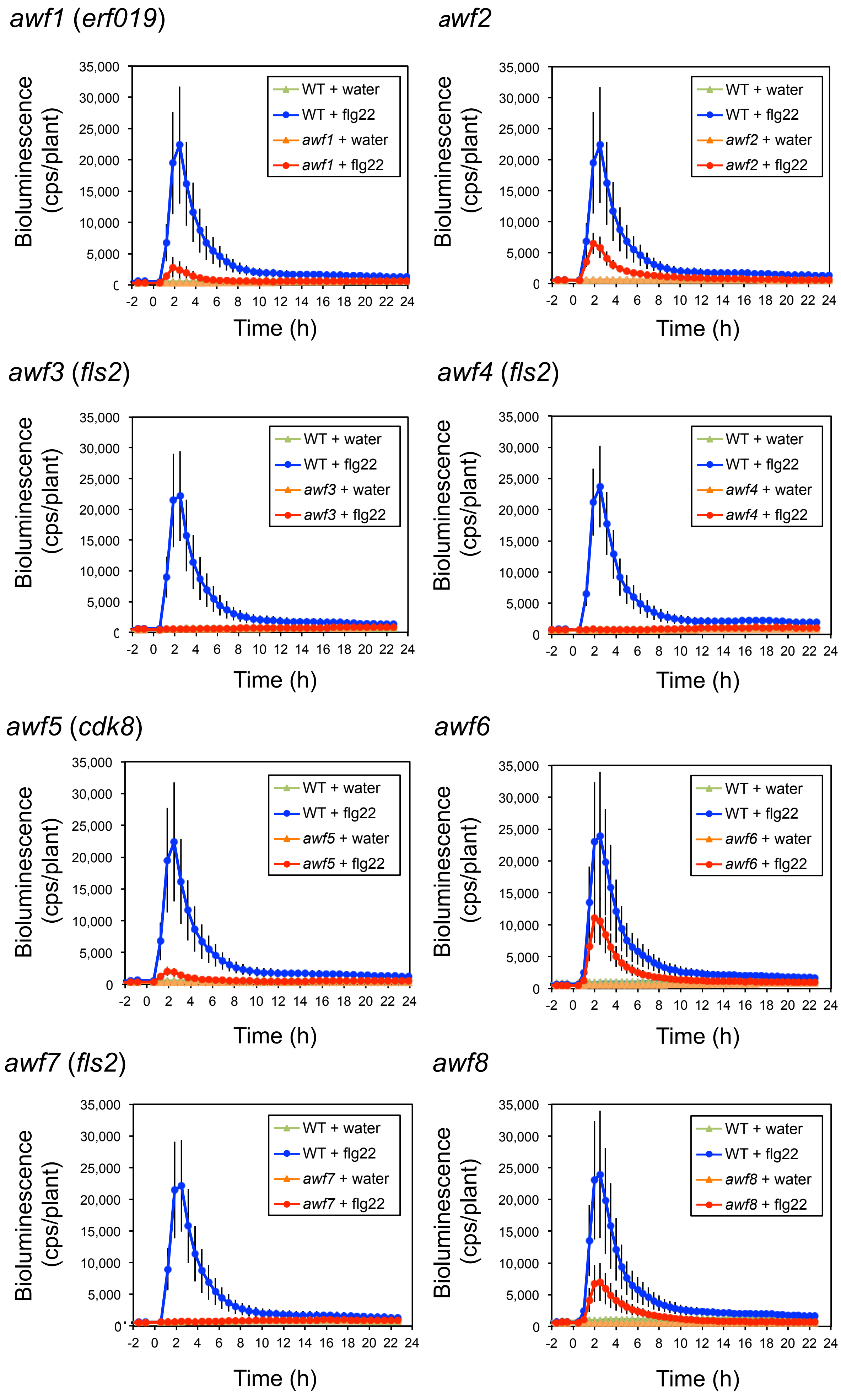
awf1 (erf019)
awf2
awf3 (fls2)
awf4 (fls2)
awf5 (cdk8)
awf6
awf7 (fls2)
awf8
Bioluminescence (cps/plant)
Time (h)
35,000
30,000
25,000
20,000
15,000
10,000
5,000
0
-2 0 2 4 6 8 10 12 14 16 18 20 22 24
WT + water
WT + flg22
awf1 + water
awf1 + flg22
awf2 + water
awf2 + flg22
awf3 + water
awf3 + flg22
awf4 + water
awf4 + flg22
awf5 + water
awf5 + flg22
awf6 + water
awf6 + flg22
awf7 + water
awf7 + flg22
awf8 + water
awf8 + flg22

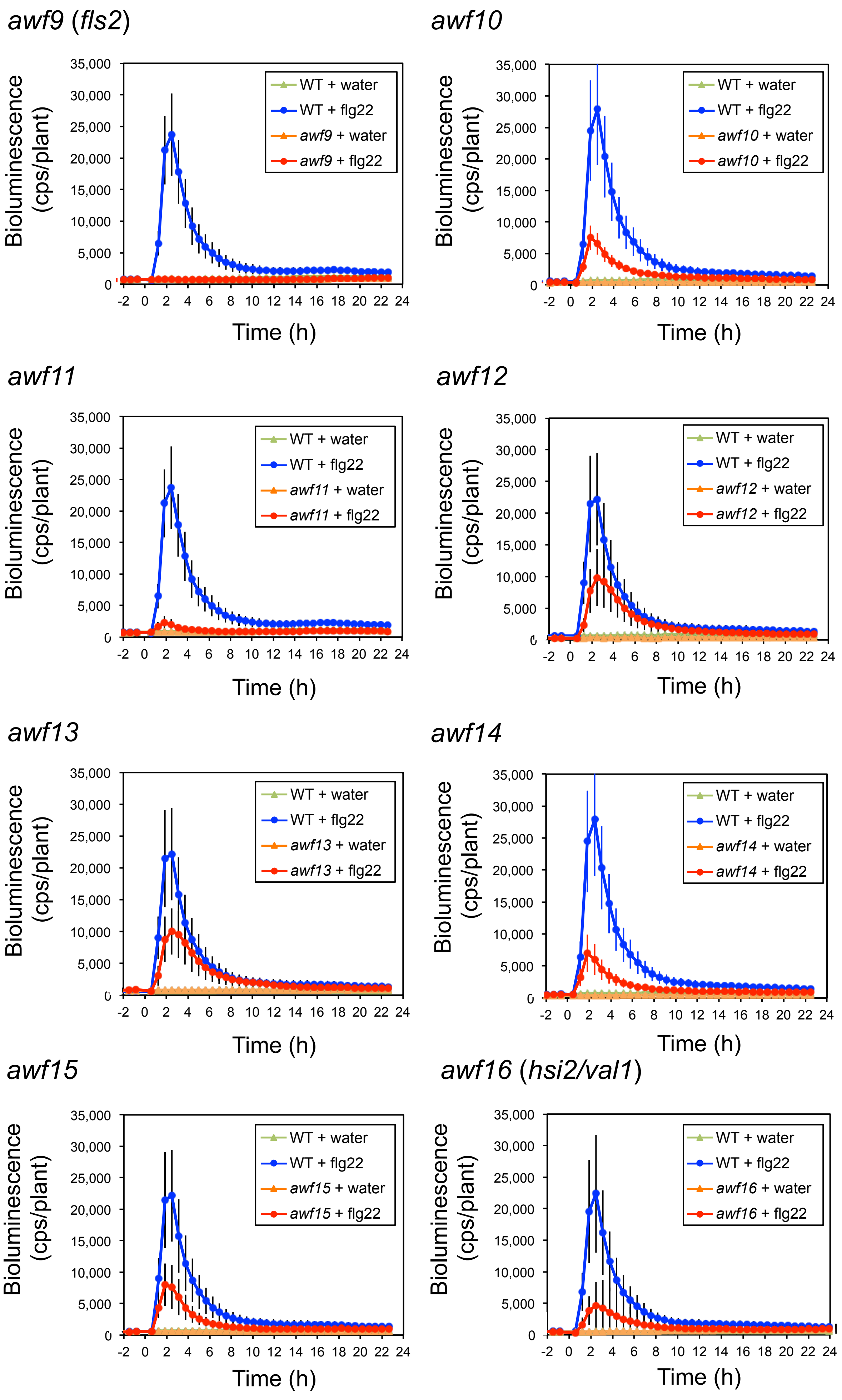

awf9 (fls2)
Bioluminescence (cps/plant)
Time (h)
WT + water
WT + flg22
awf9 + water
awf9 + flg22
awf10
Bioluminescence (cps/plant)
Time (h)
WT + water
WT + flg22
awf10 + water
awf10 + flg22
awf11
Bioluminescence (cps/plant)
Time (h)
WT + water
WT + flg22
awf11 + water
awf11 + flg22
awf12
Bioluminescence (cps/plant)
Time (h)
WT + water
WT + flg22
awf12 + water
awf12 + flg22
awf13
Bioluminescence (cps/plant)
Time (h)
WT + water
WT + flg22
awf13 + water
awf13 + flg22
awf14
Bioluminescence (cps/plant)
Time (h)
WT + water
WT + flg22
awf14 + water
awf14 + flg22
awf15
Bioluminescence (cps/plant)
Time (h)
WT + water
WT + flg22
awf15 + water
awf15 + flg22
awf16 (hsi2/val1)
Bioluminescence (cps/plant)
Time (h)
WT + water
WT + flg22
awf16 + water
awf16 + flg22

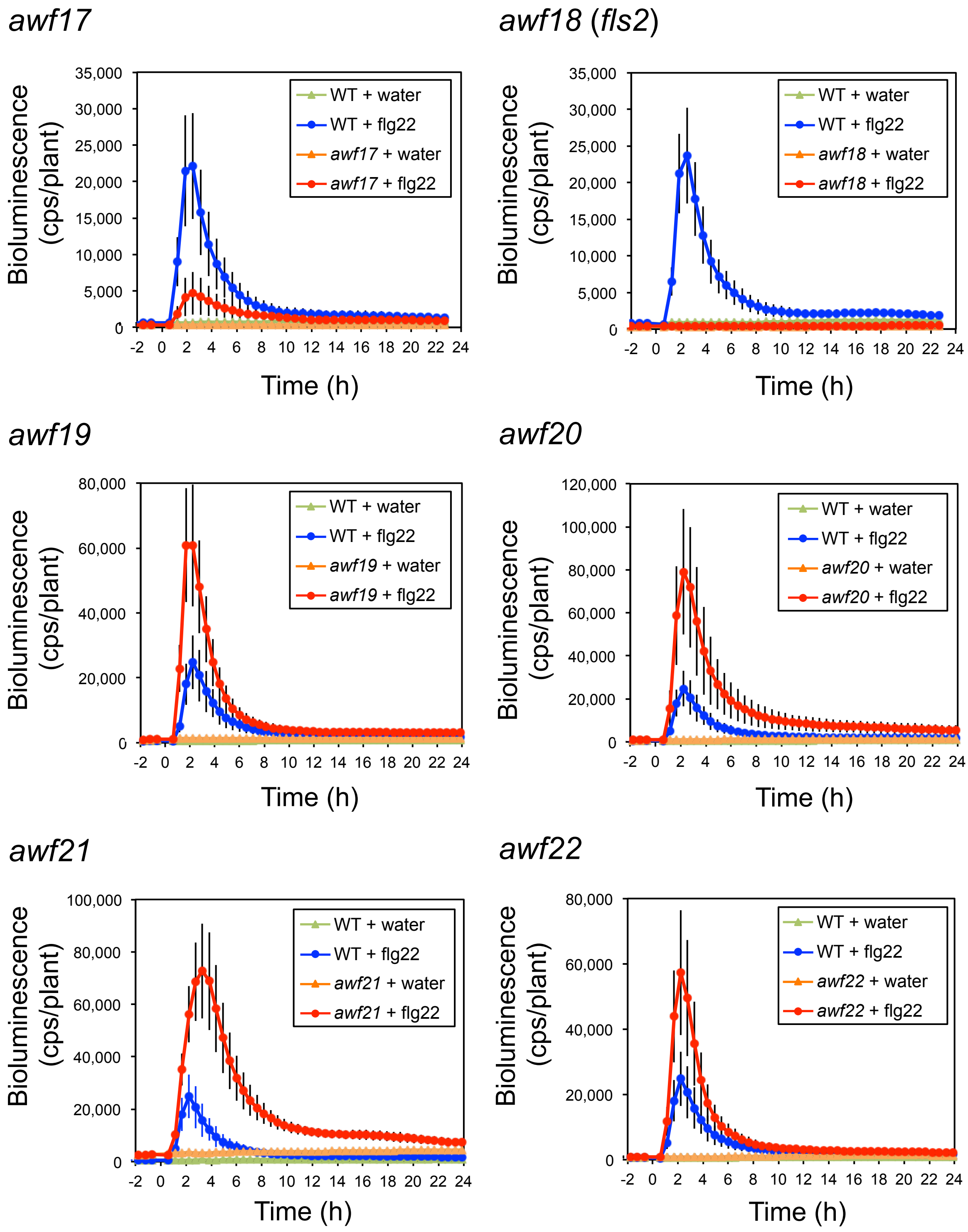


**Fig. S3** Bioluminescence phenotypes of 22 *awf* mutants as presented in Table 1. Eight day old W29-1-4 and *awf* mutants seedlings were treated with water or 0.5 μM flg22. Bioluminescence from each seedling was monitored with a real-time bioluminescence monitoring system at the indicated time points. Data are presented as mean ± SE from at least seven seedlings per treatment. Experiments were conducted three times with similar results.

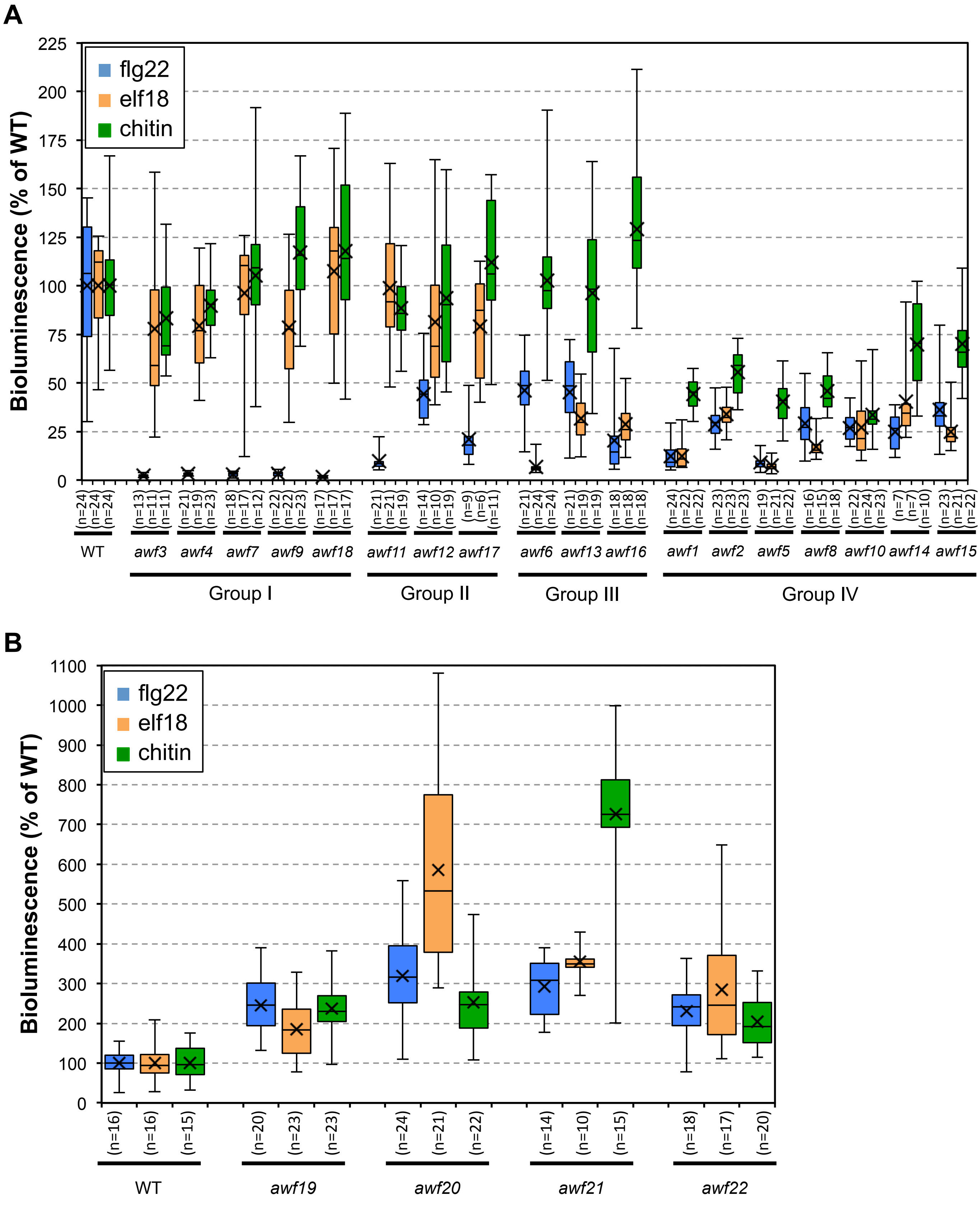


**Fig. S4** Box and whisker plot of bioluminescence phenotypes of *awf* mutants as presented in Fig. 3. **A** Low bioluminescence mutants were classified into four groups: Group I; mutants with no response to flg22, Group II; mutants showing low bioluminescence induction after flg22 treatment, Group III; mutants showing low bioluminescence induction after flg22 and elf18 treatment, Group IV; mutants showing low bioluminescence induction after flg22, elf18, and chitin treatment. **B** Mutants with increased bioluminescence induction after elicitor treatment.

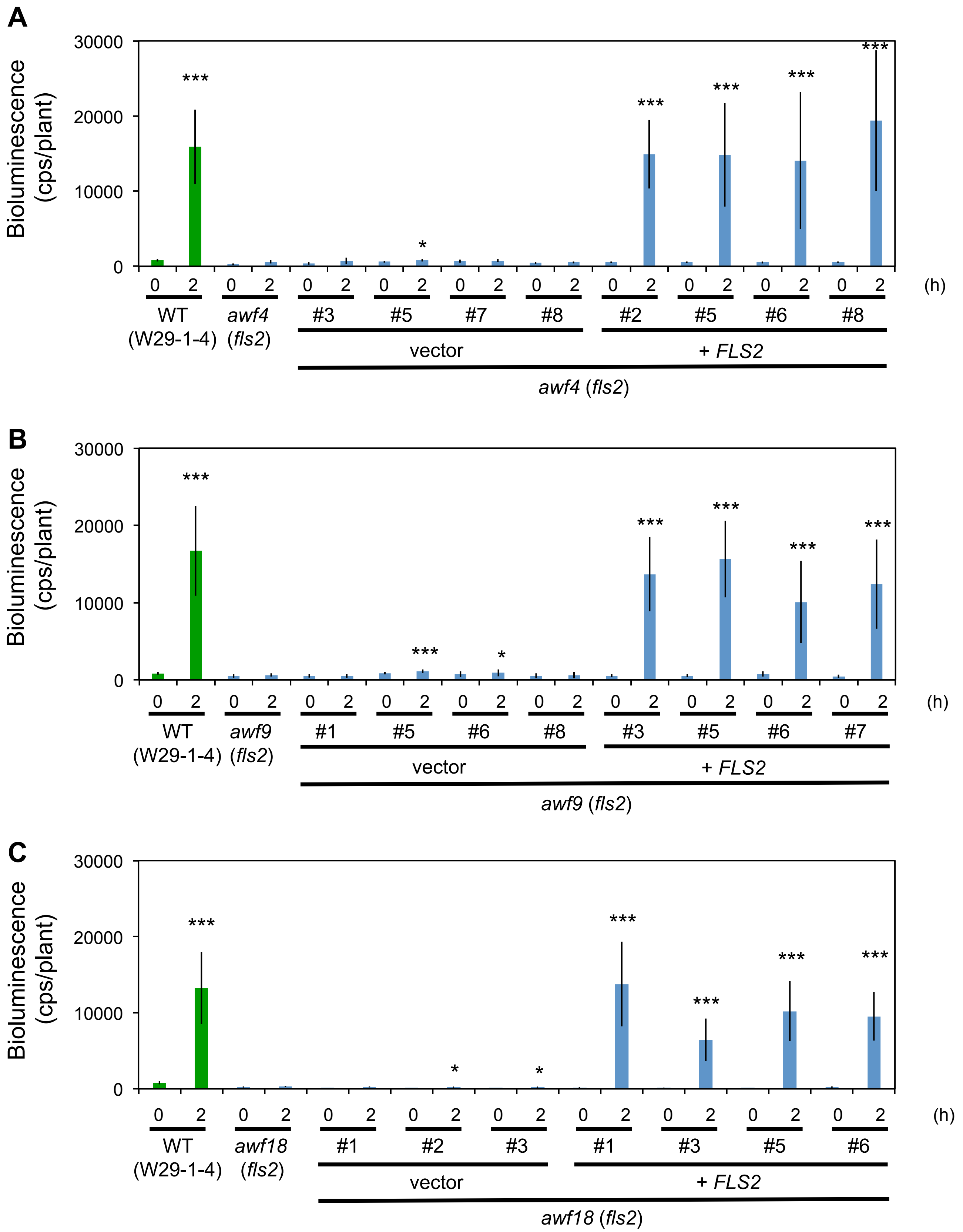


**Fig. S5** Genetic complementation of *fls2* mutants. Genetic complementation of *awf4* (**A**), *awf9* (**B**), and *awf18* (**C**) with *FLS2*. Eight day old WT (W29-1-4), *awf4*, *awf9*, and *awf18* mutants carrying the *FLS2* gene or empty vector (vector) seedlings were treated with water or 0.5 μM flg22. Bioluminescence from each seedling was monitored with a real-time bioluminescence monitoring system at the indicated time points. Data are shown as mean ± SE from at least five seedlings per treatment. Experiments were conducted twice with similar results. Asterisks indicate significant differences compared with the bioluminescence of *awf* mutants at each time point (*; $P < 0.05$, ***; $P < 0.001$, two-tailed t-tests).

WT

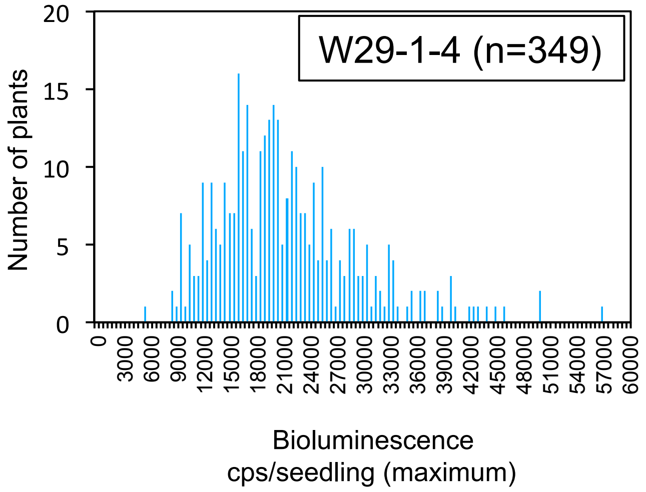
W29-1-4 (n=349)
Number of plants
0
5
10
15
20
0
3000
6000
9000
12000
15000
18000
21000
24000
27000
30000
33000
36000
39000
42000
45000
48000
51000
54000
57000
60000
Bioluminescence
cps/seedling (maximum)

Low-type

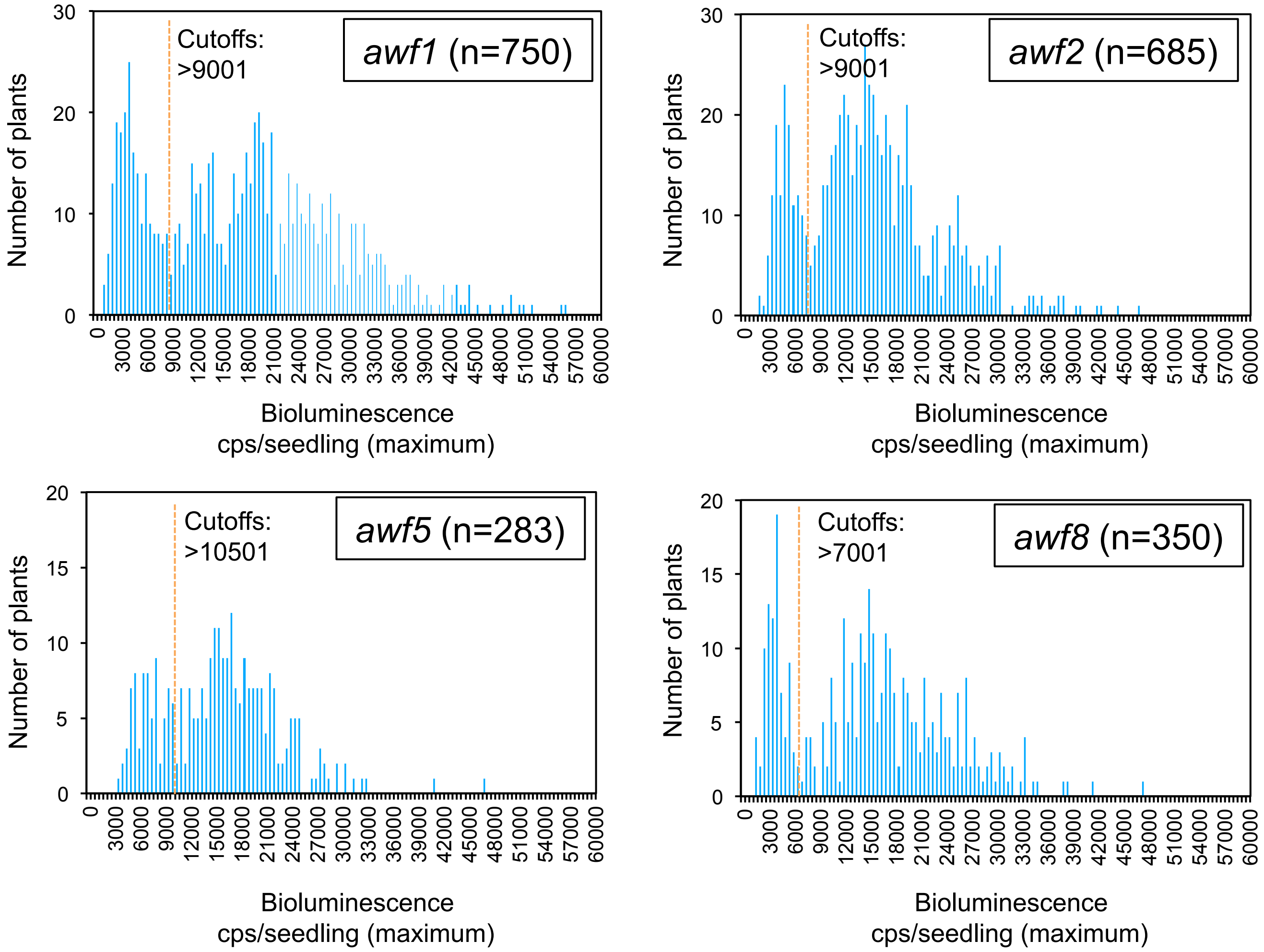
awf1 (n=750)
Cutoffs:
>9001
awf2 (n=685)
Cutoffs:
>9001
awf5 (n=283)
Cutoffs:
>10501
awf8 (n=350)
Cutoffs:
>7001
Number of plants
Bioluminescence
cps/seedling (maximum)

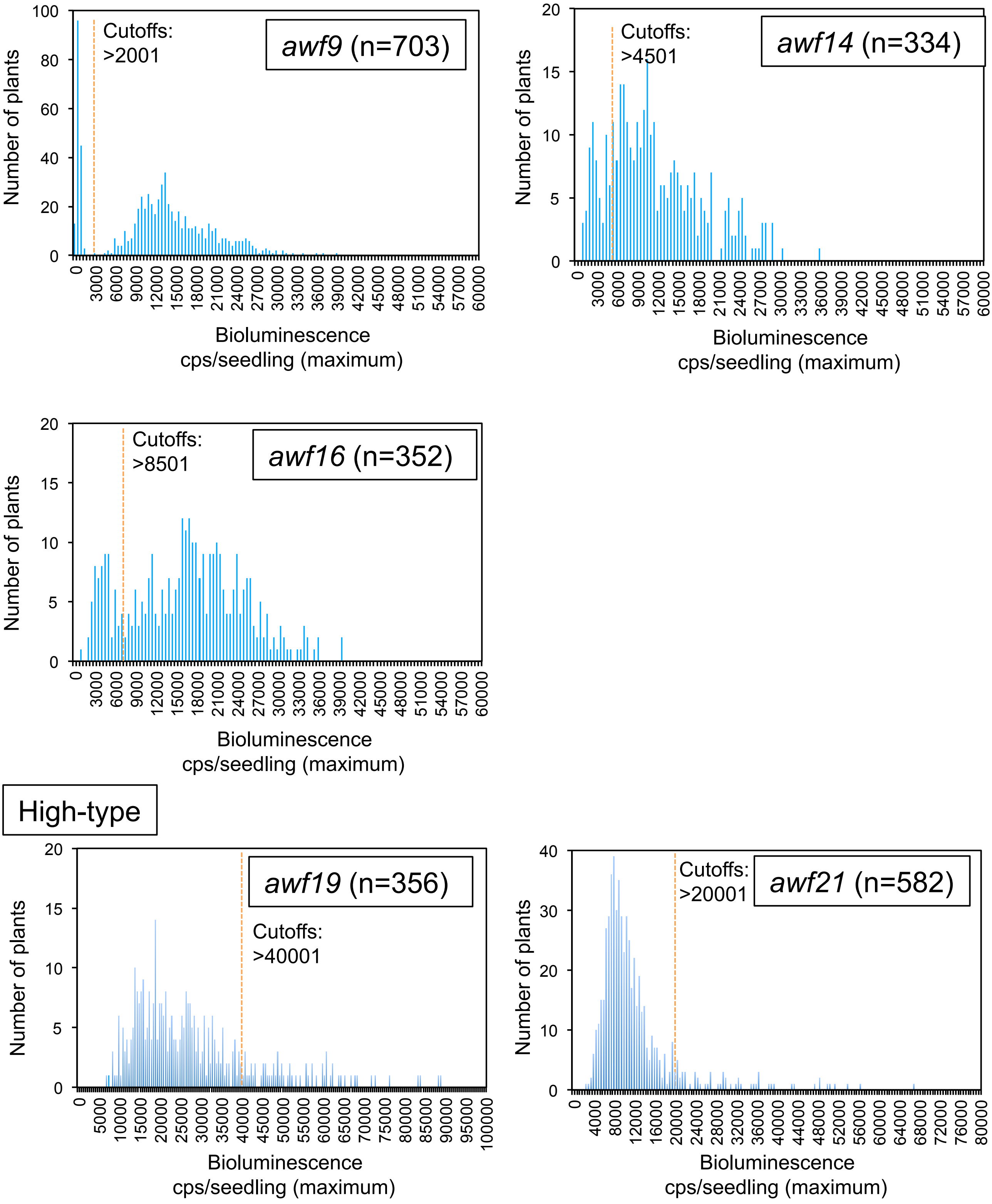


**Fig. S6** Segregation of flg22-induced bioluminescence phenotypes in $F_2$ progeny. Each panel represents the frequency distribution of flg22-induced bioluminescence (maximum value of bioluminescence, x-axis) of W29-1-4 and $F_2$ progeny derived from a cross between W29-1-4 and the designated mutant. Eight day old W29-1-4 and $F_2$ seedlings were treated with 0.5 μM flg22 and bioluminescence was monitored with a real-time bioluminescence monitoring system. Orange dashed lines indicate cutoffs used for mutant and wild-type groups.

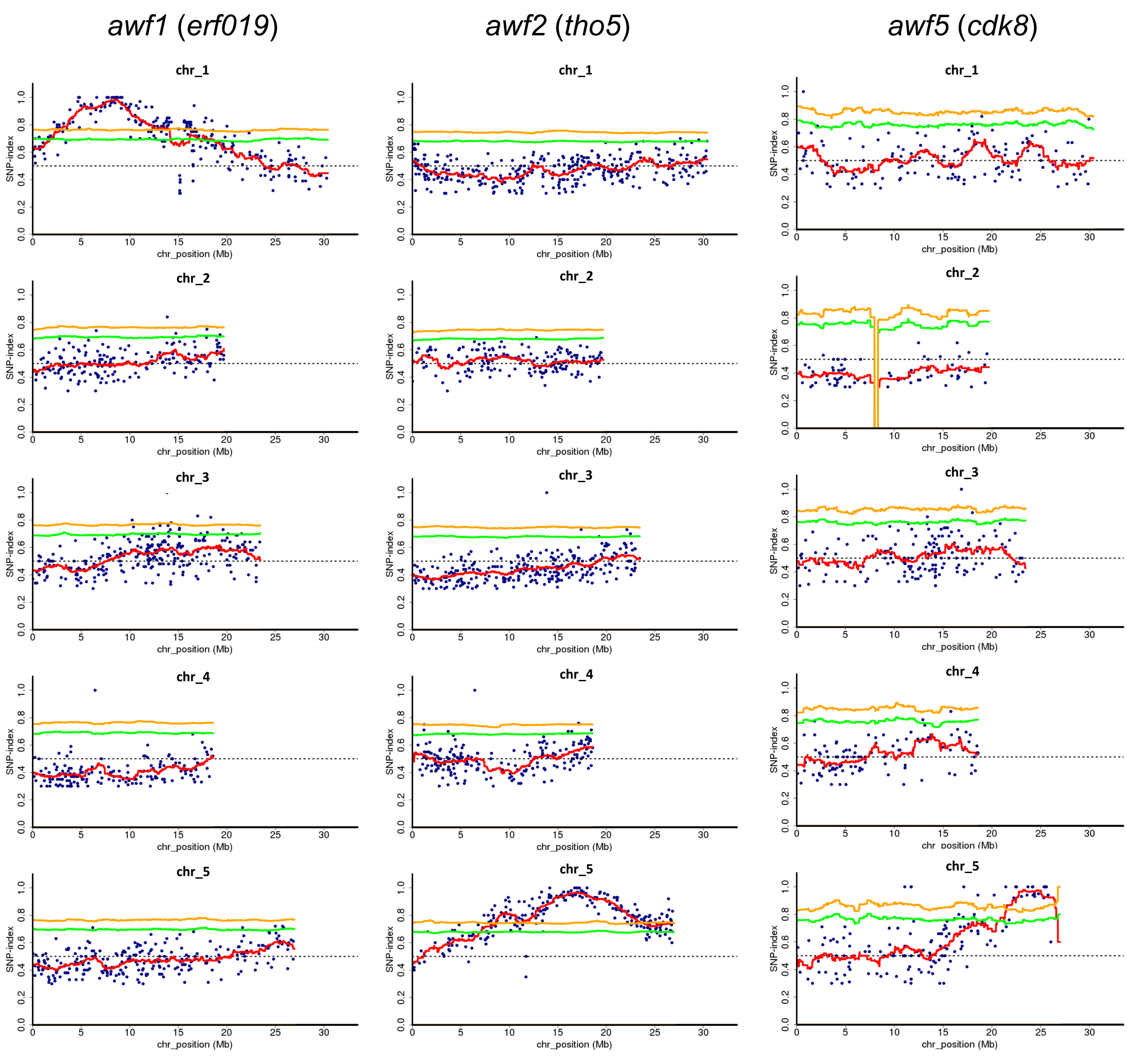
awf1 (erf019)
awf2 (tho5)
awf5 (cdk8)
chr_1
chr_2
chr_3
chr_4
chr_5
SNP-index
chr_position (Mb)

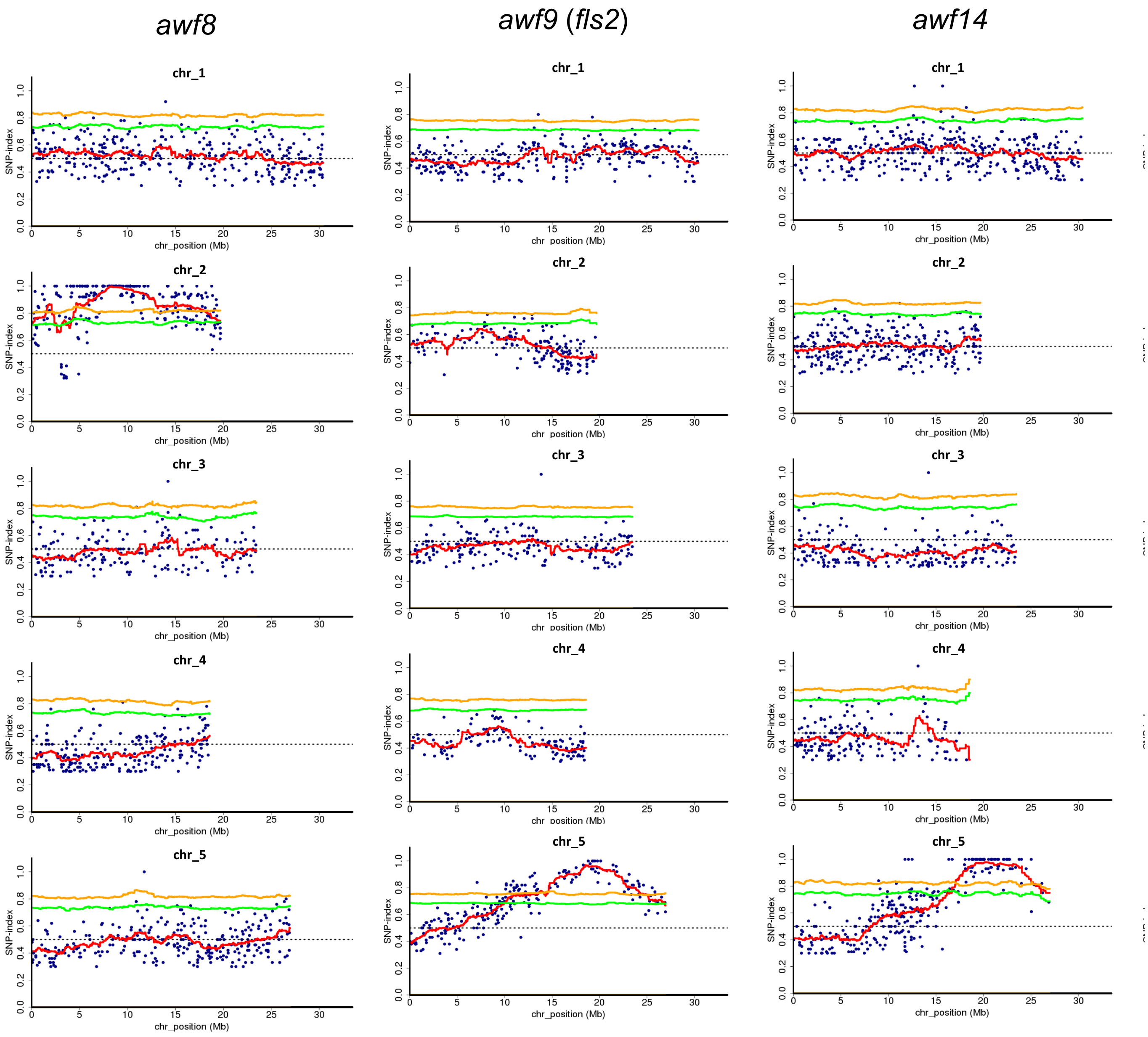

awf8
awf9 (fls2)
awf14
chr_1
chr_2
chr_3
chr_4
chr_5
SNP-index
chr_position (Mb)

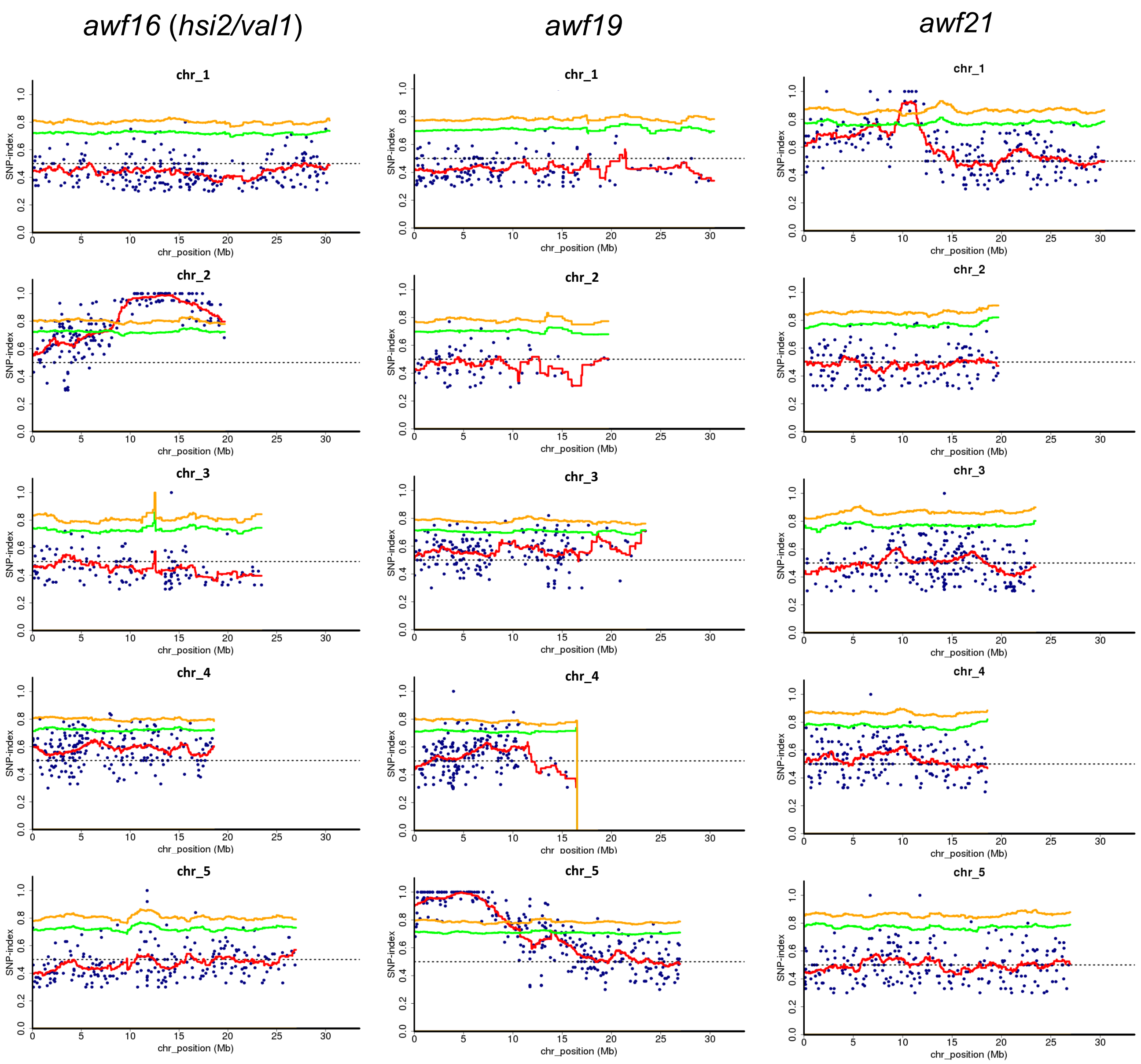


**Fig. S7** MutMap SNP-index plots of eight *awf* mutants. MutMap SNP-index plots of the five Arabidopsis chromosomes. The genomic region with the highest SNP-index peak points to the location of the causative mutation. After a cross between each mutant and the W29-1-4 line (WT), resulting $F_2$ progeny were tested for bioluminescence, and DNA from 30 $F_2$ progeny with mutant phenotypes were bulked and subjected to whole genome sequencing and MutMap analysis. Blue dots represent SNPs in the mutant. The red line represents average SNP-index values across a 2 Mb sliding window with 10 kb increments. The green and yellow lines show the 95% or 99% confidence limit, respectively, of SNP-index values under the null hypothesis of SNP-index = 0.5.

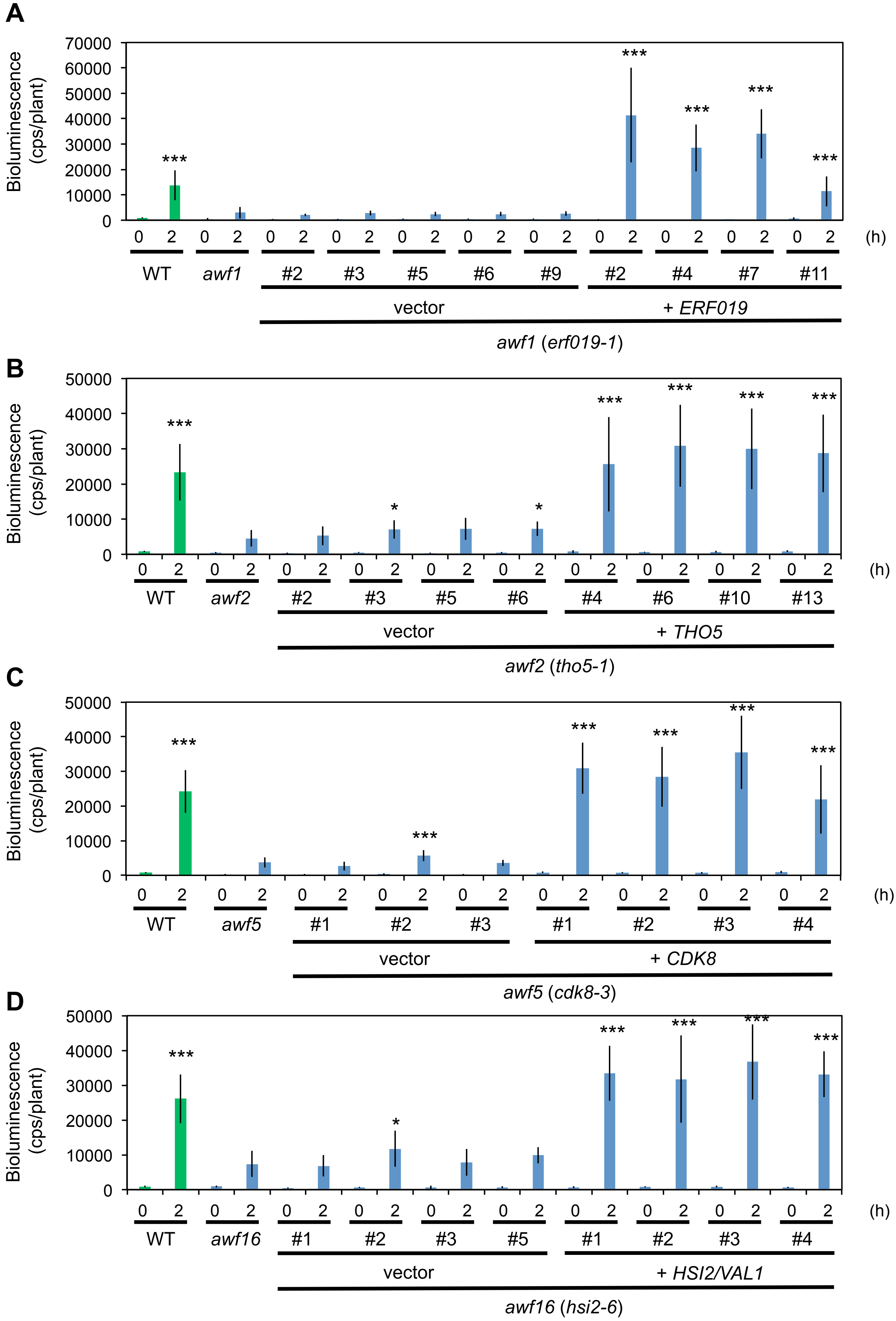
A
Bioluminescence (cps/plant)
70000
60000
50000
40000
30000
20000
10000
0
***
0 2 0 2 0 2 0 2 0 2 0 2 0 2 0 2 0 2 0 2 0 2 (h)
WT
awf1
#2 #3 #5 #6 #9 #2 #4 #7 #11
vector
+ ERF019
awf1 (erf019-1)
B
Bioluminescence (cps/plant)
50000
40000
30000
20000
10000
0
***
*
0 2 0 2 0 2 0 2 0 2 0 2 0 2 0 2 0 2 0 2 (h)
WT
awf2
#2 #3 #5 #6 #4 #6 #10 #13
vector
+ THO5
awf2 (tho5-1)
C
Bioluminescence (cps/plant)
50000
40000
30000
20000
10000
0
***
0 2 0 2 0 2 0 2 0 2 0 2 0 2 0 2 0 2 (h)
WT
awf5
#1 #2 #3 #1 #2 #3 #4
vector
+ CDK8
awf5 (cdk8-3)
D
Bioluminescence (cps/plant)
50000
40000
30000
20000
10000
0
***
*
0 2 0 2 0 2 0 2 0 2 0 2 0 2 0 2 0 2 0 2 (h)
WT
awf16
#1 #2 #3 #5 #1 #2 #3 #4
vector
+ HSI2/VAL1
awf16 (hsi2-6)

**Fig. S8** Bioluminescence of genetic complementation lines presented in Fig. 6. Genetic complementation of *awf1* with *ERF019* (**A**), *awf2* with *THO5* (**B**), *awf5* with *CDK8* (**C**), and *awf16* with *HSI2/VAL1* (**D**). Eight day old seedlings of WT (W29-1-4) the four mutants *awf1*, *awf2*, *awf5*, *awf16*, as well as mutants transformed with the indicated gene or empty vector (vector) were treated with 0.5 μM flg22 and bioluminescence was monitored with a real-time bioluminescence monitoring system. Data are shown as bioluminescence mean (0 and 2 h after treatment) ± SE from at least five seedlings per line. Asterisks indicate significant differences compared with the bioluminescence of *awf* mutants at each time point (*; $P < 0.05$, ***; $P < 0.001$, two-tailed t-tests).

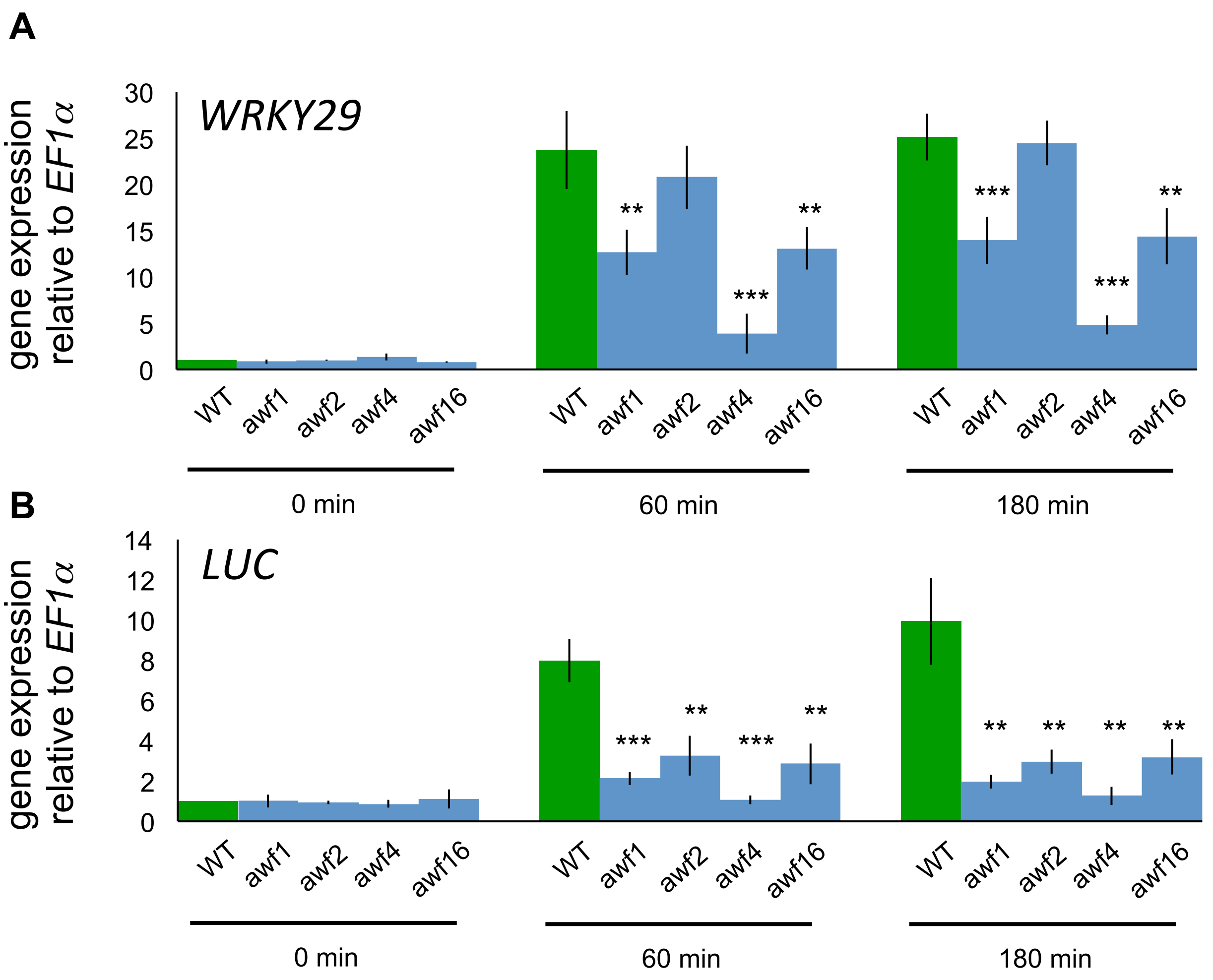


**Fig. S9** Gene expression analysis in seedlings of *awf* mutants treated with flg22. Eight day old seedlings of WT (W29-1-4), *awf1* (*erf019-1*), *awf2* (*tho5-1*), *awf4* (*fls2*), and *awf16* (*hsi2-6*) were treated with 0.5 μM flg22. Gene expression values of *WRKY29* (**A**) and *Luciferase* gene (**B**) are relative to the *EF1α* housekeeping gene (*At1g07920*) and were normalized to untreated WT seedlings. Values are shown as mean ± SE. Asterisks indicate significant differences compared with WT value at each time point (*; $P < 0.05$, **; $P < 0.01$, ***; $P < 0.001$, two-tailed t-tests). Experiments were conducted twice with similar results.

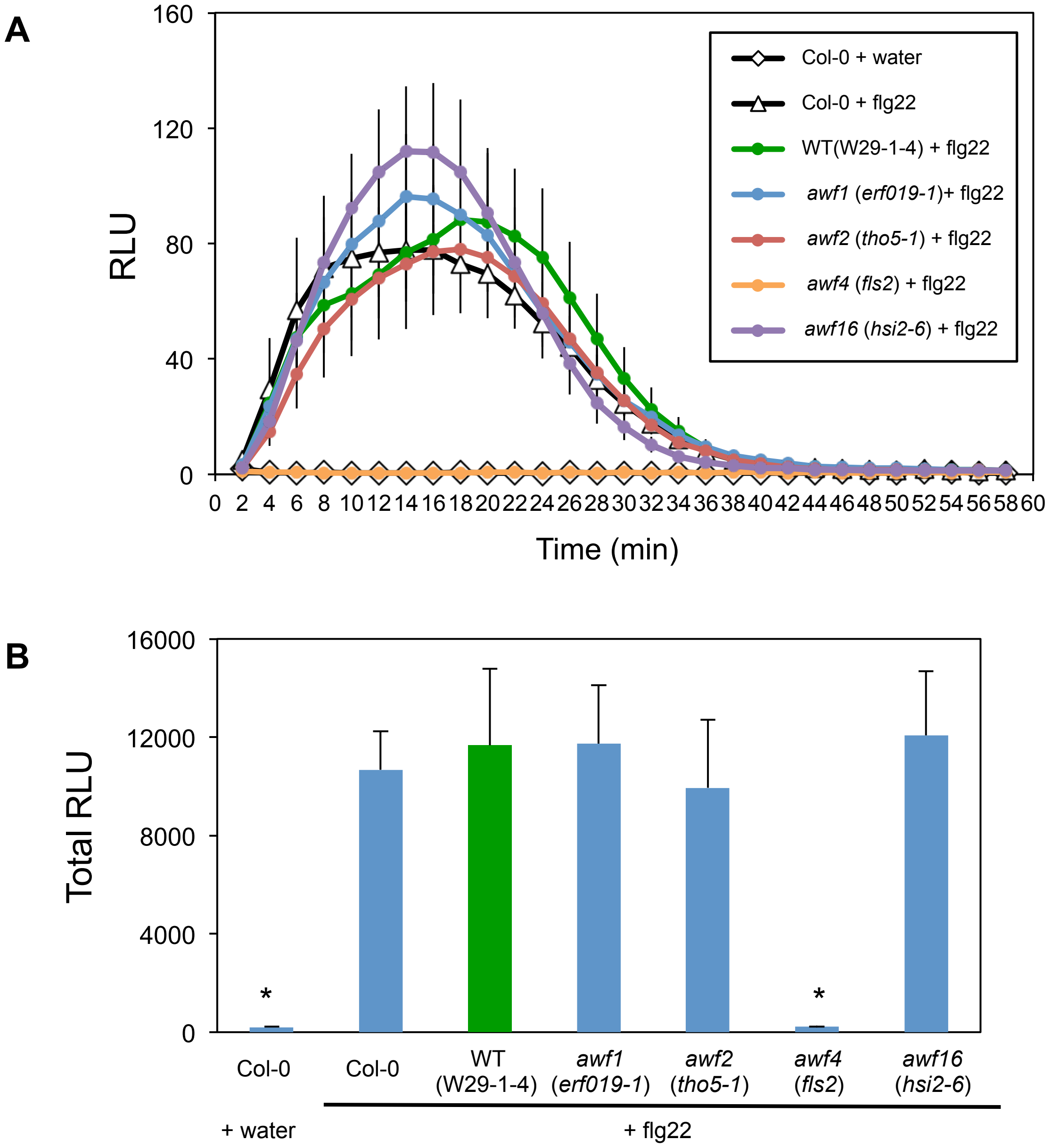


**Fig. S10** flg22-inducing early ROS production in *awf* mutants. **A** Leaf discs from five week old Col-0, WT, *awf1*, *awf2*, *awf4*, and *awf16* plants were treated with water or 0.5 μM flg22. ROS production from each leaf disc was measured in a luminol-based assay as relative light units (RLU). Data are shown as mean ± SE from 36 leaf discs per treatment derived from three independent experiments. **B** Result was shown in total RLU over the time-course. Asterisks indicate significant differences compared with WT value at each time point (*; *P* < 0.05, two-tailed t-tests).

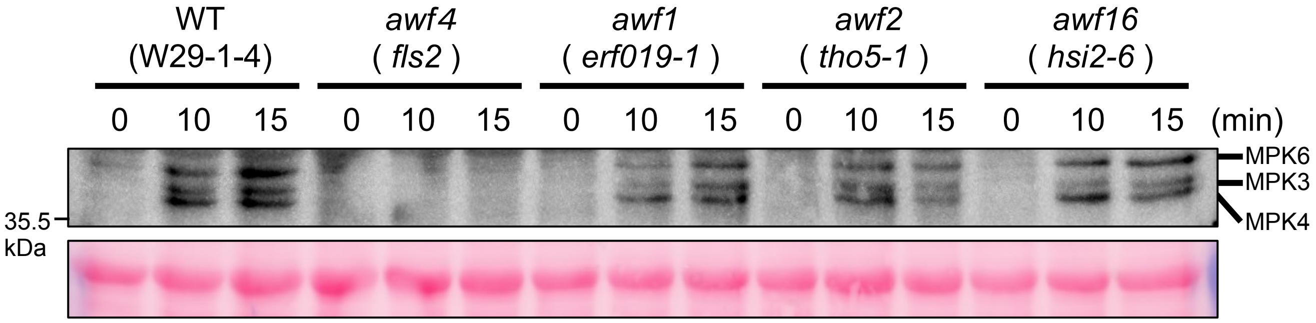


**Fig. S11** MAPK activation assay in *awf* mutants. Eight day old seedlings of WT, *awf1*, *awf2*, *awf4,* and *awf16* were treated with 0.5 μM flg22. Putative MAPKs (MPK3, MPK4, and MPK6) are indicated (upper panel). Blot were stained with Ponceau S and the protein band corresponding to the RuBisCO large subunit showed equal loading (lower panel).

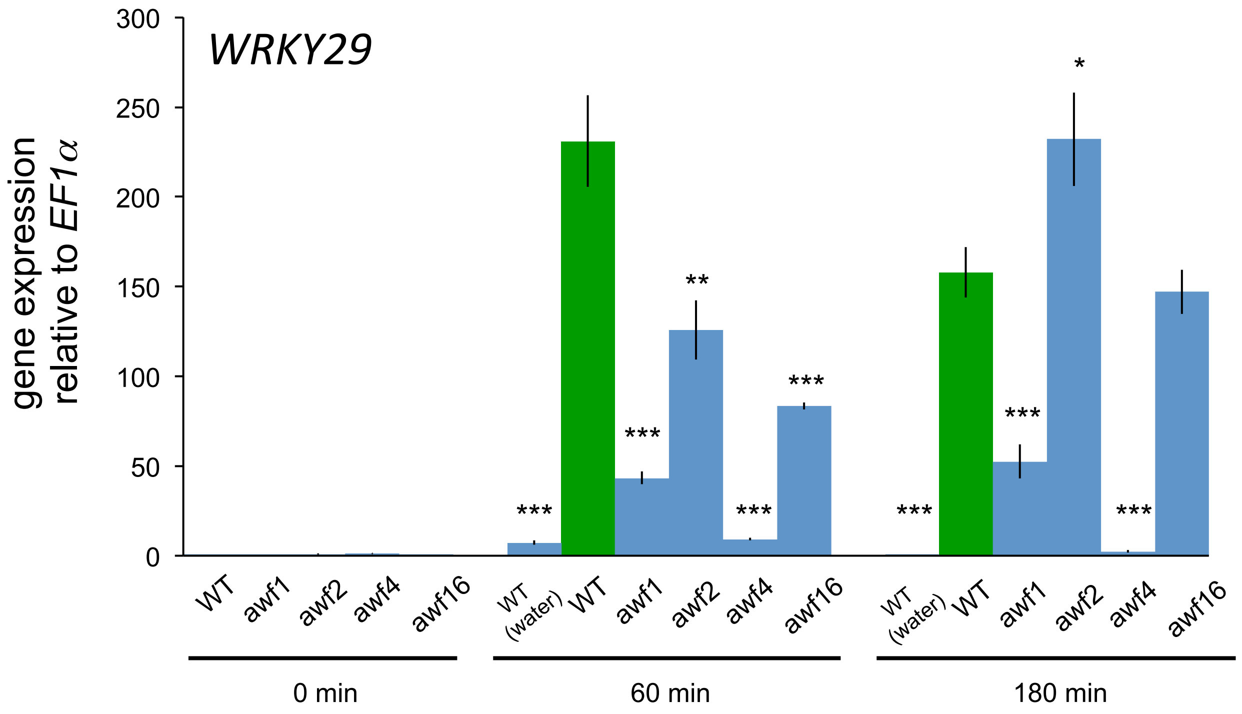


**Fig. S12** *WRKY29* expression analysis in *awf* mutant leaves treated with flg22. Five week old plants of WT (W29-1-4), *awf1* (*erf019-1*), *awf2* (*tho5-1*), *awf4* (*fls2*), and *awf16* (*hsi2-6*) were infiltrated with 0.5 μM flg22. *WRKY29* expression values are relative to the *EF1α* housekeeping gene (*At1g07920*) and were normalized to untreated WT leaves. Values are shown as mean ± SE (n=3). Asterisks indicate significant differences compared with WT value at each time point (*; $P < 0.05$, **; $P < 0.01$, ***; $P < 0.001$, two-tailed t-tests). Experiments were conducted twice with similar results.
